# Particle-Associated Prokaryotic Community Structures in the Bering and Chukchi Seas

**DOI:** 10.64898/2026.11.02.763581

**Authors:** Rei Shibue, R. Eric Collins

## Abstract

Free-living and particle-associated bacteria play distinct roles in the cycling of organic matter, but how these lifestyles are partitioned in the highly productive, rapidly warming Bering and Chukchi Seas is largely unknown. During the Arctic Shelf Growth, Advection, Respiration and Deposition Rate Experiments (ASGARD) cruise in June 2017, a period of anomalously warm sea surface temperatures, we collected 78 samples at three depths (surface, middle, and bottom) at 12 process stations and sequentially filtered them onto 20 µm, 3 µm, and 0.2 µm filters. Using 16S rRNA gene amplicon sequencing and hierarchical clustering of amplicon sequence variant (ASV) relative abundances, we resolved four sample clusters. Although pore size was not used to build the clusters, communities separated into a free-living cluster associated mainly with Anadyr Water (FL1; 0.2 µm fraction) and three particle-associated clusters (PA1–PA3; 3 and 20 µm fractions), several of which were enriched in Bacteroidia. Alphaproteobacteria, Gammaproteobacteria, and Bacteroidia were present in all size fractions, with Gammaproteobacteria most biased toward the free-living fraction. Community structure did not differ with depth, consistent with the shallow, weakly stratified shelf, and distance-based redundancy analysis identified salinity as the sole significant environmental factor distinguishing communities. These results suggest that lifestyle and water-mass origin, rather than depth, structure size-fractionated bacterial communities on the Pacific Arctic shelf, and that freshening and changes in bloom dynamics could reshape particle-associated communities in this region.

## 1 Introduction

Microbial communities are central to marine ecosystems, contributing to primary production and organic-matter decomposition and driving biogeochemical cycles (Azam & Malfatti, 2007; Falkowski et al., 2008). As primary producers they channel carbon and energy to higher trophic levels, and by remineralizing organic matter they cycle key elements such as carbon, nitrogen, and sulfur; through the microbial carbon pump they further convert labile organic matter into long-lived recalcitrant dissolved organic carbon (Jiao et al., 2024; Legendre & Le Fèvre, 1995).

The Arctic Ocean is a geographically unique setting for these processes. Although it represents only about 3 % of the global ocean by area and roughly 1 % by volume, it holds about 20 % of the world’s continental shelf and receives close to 11 % of global riverine input (Carmack, 2007; Tomczak & Godfrey, 1994). It is largely enclosed by land, seasonally ice-covered, and generally nitrogen-limited, with primary production dominated by picoeukaryotic algae rather than cyanobacteria such as *Synechococcus* (Pedrós-Alió et al., 2015; Simpson et al., 2008).

The region is changing rapidly. Sea-ice extent and thickness are declining, and Arctic amplification has warmed the region roughly four times faster than the global ocean (Rantanen et al., 2022; Serreze & Barry, 2011). Warming is reinforced by heat advected from the Atlantic (Atlantification) and the Pacific, alongside acidification, freshening, and stronger stratification that suppress the vertical supply of nutrients (Polyakov et al., 2020; Tremblay & Gagnon, 2009). These changes reach the microbial communities as well. For instance, their composition shifted after the record loss of multi-year sea ice in 2007 (Comeau et al., 2011). Net primary production rose by about 57 % from 1998 to 2018 (Lewis et al., 2020). New harmful algal blooms have also appeared, including the toxic *Alexandrium catenella* bloom in the Chukchi Sea in 2022 (Fachon et al., 2025). Because they respond to drastic environmental change quickly, microbial communities serve as sensitive early indicators of Arctic change.

This study focuses on the Bering and Chukchi Seas, the Pacific gateway to the Arctic Ocean and among the most productive regions on Earth (Grebmeier, 2012; Springer et al., 1996). The shallow (∼50 m) Bering–Chukchi shelf receives heat and nutrients from Pacific inflow that fuel seasonal phytoplankton blooms whose timing depends strongly on sea-ice cover and water-column stratification (Hunt et al., 2002; Woodgate et al., 2005). Nutrient-rich Pacific Winter Water sustains under-ice production, and net primary production in the Chukchi Sea has nearly doubled since the late 1990s (Arrigo et al., 2014; Lewis et al., 2020). The region’s waters are classically divided into Alaskan Coastal, Bering Shelf, and Anadyr waters (Coachman et al., 1975), recently refined into an all-season scheme of warm and cool coastal and shelf water masses (Danielson et al., 2020).

Marine bacterial communities can be broadly divided into two microbial lifestyles: free-living (FL) cells suspended in the water column and particle-associated (PA) cells attached to suspended or sinking particles. The FL and PA fractions of marine bacterial communities have been compared from estuaries and coastal waters to the open sea (Crespo et al., 2013; Hollibaugh et al., 2000; LaMontagne & Holden, 2003; Rieck et al., 2015; Roth Rosenberg et al., 2021; Selje & Simon, 2003; Zhang et al., 2007). These studies have generally shown that FL and PA communities are taxonomically distinct, with many taxa restricted to a single fraction. For example, only about a quarter of the operational taxonomic units (OTUs; proxies for bacterial species) were shared between fractions in the northwestern Mediterranean (Crespo et al., 2013), and more than half were PA-exclusive in the Baltic Sea (Rieck et al., 2015). A recurrent pattern has emerged in which FL communities are dominated by Alphaproteobacteria, whereas PA communities are enriched in Gammaproteobacteria and Bacteroidetes (Crespo et al., 2013; Zhang et al., 2007). These temperate studies establish the FL–PA distinction as an ecologically meaningful but environmentally influenced feature of marine bacterial communities.

In the Arctic, FL and PA fractions of bacterial communities have been examined mainly in fjords, on productive shelves, and in river-influenced seas (Jain & Krishnan, 2017; Jain et al., 2019, 2021; Kellogg & Deming, 2009; Ortega-Retuerta et al., 2013; Sperling et al., 2013). As in temperate systems, FL and PA communities are generally distinct, especially in open, offshore waters, and tend to show the same taxonomic contrast. Bacteroidetes and Gammaproteobacteria are enriched in the PA fraction, whereas Alphaproteobacteria dominate the FL fraction (Ortega-Retuerta et al., 2013). PA fractions also tend to harbor higher diversity (Jain et al., 2019). However, the strength of this separation is not uniform and is closely tied to the distinctive hydrography of the Arctic. Glacial meltwater, terrestrial runoff, and water mass gradients can act on the two fractions differently, so that FL communities often vary more among locations while PA communities remain comparatively similar (Jain & Krishnan, 2017; Ortega-Retuerta et al., 2013). More specifically, the biochemical quality of particles can structure PA communities and favor polymer-degrading taxa such as Bacteroidetes and Verrucomicrobia (Jain et al., 2019). This is especially true of their carbohydrate content. The two lifestyles are increasingly regarded as spatially connected assemblages shaped by both dispersal and local environmental selection rather than as fixed habitats (Jain et al., 2021). Together, these studies show that the FL–PA distinction persists in cold, seasonally productive, and freshwater-influenced polar waters, but that its expression is strongly governed by water masses and particle supply.

FL and PA communities in the Bering and Chukchi region have also been studied. For example, Qin et al. (2025) compared suspended and sinking particles at ten western-Arctic stations including the Bering and Chukchi regions. They found that eukaryotic communities differed between particle types, while prokaryotic communities differed only slightly, at the level of individual taxa, and carbon sinking flux increased with chlorophyll a and diatom-associated taxa. In the same region, the study extends the FL/PA perspective to sinking particles, suggesting a pathway by which diatom blooms may drive carbon export. However, despite such studies, how bacterial communities in this region are structured in response to algal blooms remains only partly understood. Given the intense algal blooms and high primary productivity that characterize these seas, further work is needed to understand how bloom dynamics shape the structure of local bacterial communities. Here, using 16S rRNA gene amplicon sequencing of 78 samples collected at three depths (surface, middle, and bottom) across 12 stations during the Arctic Shelf Growth, Advection, Respiration and Deposition Rate Experiments (ASGARD) 2017 cruise, we aim to (i) characterize the composition and spatial structure of bacterial communities in the Bering and Chukchi Seas, (ii) examine how the diatom bloom influenced community composition, (iii) identify relationships between the communities and physicochemical parameters of water masses from which they derived, (iv) assess how physicochemical changes associated with ongoing warming may affect these communities, and (v) compare free-living and particle-associated communities to determine whether the environmental drivers of change differ by fraction.

This study links community structure to bloom dynamics and to physicochemical properties of water masses. Through this approach, it provides a baseline for understanding how microbial communities in this region respond to environmental change, and how different lifestyles (free-living vs. particle-associated) shape microbial communities.

## 2 Methods

### 2.1 Sample collection

Seawater was sampled during the ASGARD project between 9 and 28 June 2017 aboard the R/V Sikuliaq. Over the ASGARD 2017 cruise, 12 process stations were occupied, and four casts were carried out for sensor profiling, nutrient analysis, and zooplankton incubation at those stations. The ASGARD 2017 cruise visited 68 unique stations, many of which were associated with the Distributed Biological Observatory (DBO), including the DBO2 line in the northern Bering Sea and the DBO3 line in the southern Chukchi Sea. Samples were collected from surface, middle (pycnocline) and bottom depths between 2.5 m (surface) and 55 m (bottom), with modal depths per station at 5 m, 20 m, and 40 m. Seawater was sampled from Niskin bottles on a rosette with an attached Sea-Bird CTD (Sea-Bird Electronics Inc., Bellevue, WA, USA).

Samples were collected in two phases, here labeled “survey” and “process” stations. The survey stations (62 total) were relatively short stops where only the CTD rosette and zooplankton nets were deployed; process stations (12 total) were full-day stops, including four CTD casts conducted for sensor profiling, nutrient analysis, zooplankton incubations, and more. Some stations were visited during both phases. Figure 1 shows the process stations of ASGARD 2017 and the near-surface hydrography during the cruise obtained by the flow-through thermosalinograph. The average near-surface temperature was 4.9 °C (range 1.1–10.9 °C), and salinity 31.6 (range 25.0–32.9). Six stations (CBE1, CBW5, CL1, CL3, DBO3.3, and IL) were sampled in both the survey and process phases. All geographic maps were produced in R with the ggmap package (Kahle & Wickham, 2013). Basemap data are © Stadia Maps, © Stamen Design, © OpenMapTiles and © OpenStreetMap contributors.

**Figure 1.**
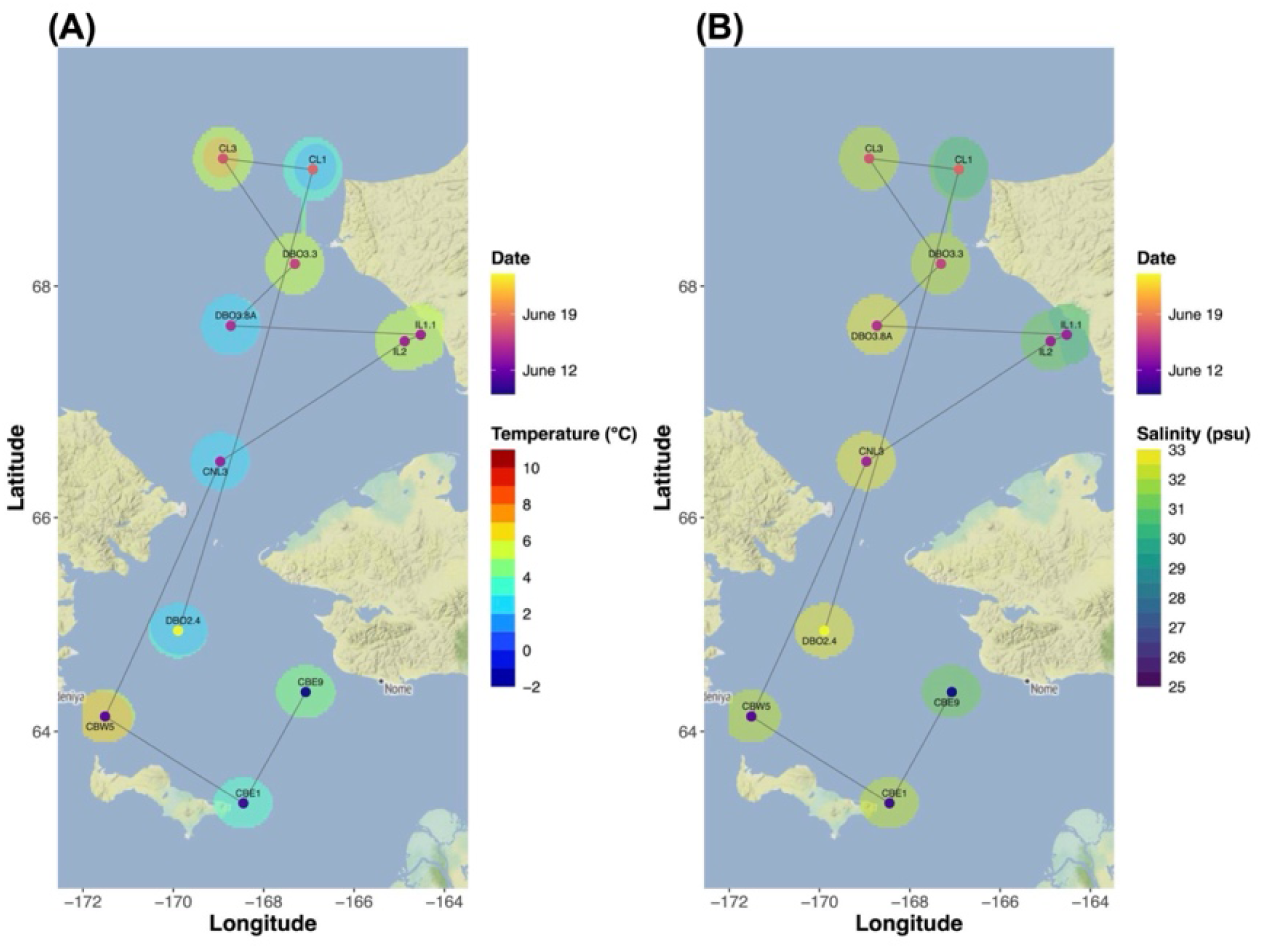
Process Stations and Near-surface Hydrography During the ASGARD 2017 Cruise. Process stations of the ASGARD 2017 cruise across the Bering and Chukchi Seas. Each station is colored by its sampling timing, is connected with the grey line in chronological order to indicate the cruise track, and is labeled by its station name based on pre-determined lines: CBE (Chirikov Basin East transect), CBW (Chirikov Basin West transect), DBO (Distributed Biological Observatory), CNL (Convention Line transect), IL (Isilivik Lagoon transect), and CL (Cape Lisburne transect). The filled background shows the near-surface (A) temperature and (B) salinity, interpolated from the station values by inverse-distance weighting and masked to within 30 km of a station.

### 2.2 Sample filtration

Seawater samples were transferred from Niskin bottles to 20 L cubitainers and stored at 4 °C until filtration within six hours. Using a peristaltic pump, 1–5 L of seawater per depth was filtered. Samples were collected with sequential filtration through a 47 mm-diameter 20 µm-pore size nylon net filter, then a 47 mm-diameter 3 µm-pore size membrane filter, and finally through a 0.2 µm-pore size Sterivex cartridge filter (all filters from Millipore Sigma, Burlington, MA, USA). Filters were stored at −20 °C until processing.

Nutrient samples were drawn with 60 mL syringes immediately after CTD recovery and filtered through 0.45 µm Nuclepore filters, and kept frozen below −20 °C until processed colorimetrically by autoanalyzer post-cruise (Gordon, Jennings and Krest 1993). Nitrate, phosphate and silicate concentrations were provided by S. Danielson (University of Alaska Fairbanks, unpublished data).

### 2.3 DNA extraction, amplification, and sequencing

DNA was extracted from filters using the DNeasy PowerWater kit (Qiagen, Hilden, Germany) following the manufacturer’s instructions. PCR-amplification of 16S rRNA genes was used for analysis of prokaryotic communities. The KAPA HiFi HotStart PCR Kit (Kapa Biosystems, Wilmington, MA, USA) was used for PCR-amplification of 16S rRNA genes using the following conditions: one cycle at 98 °C for 1 min, 26 cycles at 98 °C for 30 s, 55 °C for 30 s, 72 °C for 30 s, and one cycle at 72 °C for five minutes. Earth Microbiome Project primers (Thompson et al., 2017) were used to target the V4 hypervariable region of 16S SSU rRNA: forward primer 515F-revised (Parada et al., 2016) (5’-GTGYCAGCMGCCGCGGTAA-3’) and reverse primer 806RB-revised (Apprill et al., 2015) (5’-GGACTACNVGGGTWTCTAAT-3’). Amplified DNA was dual-indexed using unique adapters (Glenn et al., 2016) before TruSeq library preparation and sequencing on an Illumina MiSeq in the DNA Core Lab at the University of Alaska, Fairbanks (Fairbanks, USA).

### 2.4 Analysis of 16S rRNA gene sequencing

Sequences were demultiplexed and primers were removed with cutadapt v2.8 (Martin, 2014). The amplicon sequencing data was analyzed for amplicon sequence variants (ASVs) using DADA2 in R (Divisive Amplicon Denoising Algorithm 2; Callahan et al., 2016), performing quality control, error correction, merging, chimera checks and taxonomic classification. The SILVA database v132 (Quast et al., 2012) was used to classify ASVs.

Altogether, the 16S rRNA gene comprised 3,076 prokaryotic ASVs in total. Of these 3,076 ASVs, 374 were detected (> 2 reads) in each dataset in the 78 process samples that retained *≥* 5,000 reads. Read counts were converted to relative abundances within each sample, and an ASV was retained only when its maximum relative abundance across samples exceeded 0.001 (0.1 %) and it was present in more than two samples. This filtering step left 221 ASVs from the 78 process samples.

The analyses described in the following sections were performed in R (version 4.5.2). All R scripts and outputs are publicly available at https://github.com/ReiSHIBUE/ASGARD2017_analysis.

### 2.5 Hierarchical Clustering

For the beta-diversity analysis, Bray–Curtis dissimilarities were derived from the fourth-root-transformed relative-abundance matrix with the vegan package in R (Oksanen et al., 2024). Samples were then clustered by Ward’s minimum-variance method and displayed as a heatmap built with the R packages vegan, fastcluster (Müllner, 2013), and gplots (Warnes et al., 2009). For the process dataset, Bray–Curtis dissimilarities among samples were computed from the fourth-root-transformed ASV relative-abundance matrix (221 ASVs across 78 process samples) and the samples were grouped using Ward’s method. Cutting the dendrogram at k = 4 yielded four clusters. The clustering was based on ASV composition alone; filter pore size was not used as an input. The first split of the dendrogram nevertheless separated samples collected on 0.2 µm filters from those collected on 3 and 20 µm filters. On this basis the four clusters were labelled as one free-living cluster (FL1) and three particle-associated clusters (PA1–PA3). These labels are operational, describing the size fraction in which each community was recovered rather than a directly observed lifestyle. This division was also examined through a top-down tree walk, carried out differently from the hierarchical clustering used for the survey dataset. Here, environmental variables were log-transformed (nutrients), scaled, and reduced to principal components explaining *≥* 90 % of cumulative variance; at each internal node with *≥* 5 samples per child, PC scores were compared with Wilcoxon rank-sum tests and adjusted across PCs with the Benjamini–Hochberg procedure (significance at BH-adjusted p < 0.05).

Bubble plots were generated to visualize relative abundances of ASVs belonging to three major bacterial classes (Alphaproteobacteria, Gammaproteobacteria, and Bacteroidia). For each sample, ASV relative abundances were summed to the genus level and averaged across the samples within each cluster, giving a mean relative abundance (%) per genus per cluster.

ASV richness, defined as the number of ASVs present (relative abundance > 0) in the filtered clustering set (221 ASVs), was compared among depths using Kruskal–Wallis rank-sum tests. The tests were run overall and separately within each community grouping (one free-living cluster and three particle-associated clusters), and p-values were Bonferroni-corrected for the number of within-group tests.

Ternary plots were produced in R using the Ternary package; normalized mean relative abundances of each ASV in each filter fraction were used as input (0.2 µm: n = 23 samples, 3 µm: n = 27, 20 µm: n = 28).

### 2.6 Water Mass Characterization

A temperature–salinity (T-S) diagram was constructed to characterize the water masses sampled and to visualize their relationship with the microbial community clusters. Potential density isopycnals were overlaid for reference, and water mass boundaries followed the classification of Danielson et al. (2020), reproduced in Table 1. All plots were produced with ggplot2 in R (Wickham, 2016). Seawater density isolines were computed from the EOS-80 equation of state. The correlation between each cluster and water mass was assessed by Fisher’s exact test (9,999 Monte Carlo replicates), while the significance of correlations between each cluster and depth was tested with a Kruskal–Wallis test.

**Table 1.** Water Mass Definitions in the Northern Bering and Chukchi Seas, Modified From Danielson et al. (2020)

| Water mass | Abbreviation | Temperature range (°C) | Salinity range |
|---|---|---|---|
| Anadyr Water | AnW | 0 < T < 3 | 32.5 < S < 33.8 |
| Ice Melt Water & cool Coastal Water | IMW, cCW | −2 < T < 3 | 22 < S < 30.8 |
| cool Shelf Water | cSW | 0 < T < 3 | 30.8 < S < 32.5 |
| warm Coastal Water | wCW | 3 < T < 14 | 18 < S < 30.8 |
| warm Shelf Water | wSW | 3 < T < 14 | 30.8 < S < 33.4 |
| Modified Winter Water | MWW | −1 < T < 0 | 30.8 < S < 33.8 |
| Winter Water | WW | −2 < T < −1 | 30.8 < S < 35 |

*Note.* Atlantic Water and Bering Basin Water (AtlW and BBW) are not included in this table because no samples in this study fell within their temperature–salinity ranges. Temperature (T) is in °C and salinity (S) is unitless (PSU).

Samples were assigned to the Bering or Chukchi by latitude (south or north of the Bering Strait at 65.7 °N, respectively), and potential density was calculated from temperature and salinity using the EOS-80 equation of state at surface pressure.

### 2.7 Environmental Characterization of Sample Clusters

Boxplots were generated for each measured environmental variable (temperature, salinity, dissolved oxygen, nitrate, phosphate, ammonium, chlorophyll fluorescence, chlorophyll a, sampling depth, and latitude) to explore how environmental conditions varied among microbial community clusters. Each cluster was visualized with median, interquartile range, and overlaid individual sample points using ggplot2 in R.

To classify each microbial community cluster by its bloom stage, principal component analysis (PCA) was performed on nine standardized environmental variables (with log(1+x)-transformation of right-skewed nutrient and fluorescence variables). PC1 was observed to capture the dominant gradient between pre- and post-bloom conditions, and was rescaled to the [0, 1] interval and interpreted as a “bloom index” (0 = pre-bloom, 1 = post-bloom). The index is derived from environmental data alone and uses no microbial information, so the assignment of bloom stages to clusters is independent of the clustering itself. The distribution of the bloom index across clusters was then used to assign a representative bloom stage to each cluster.

Parameter selection was conducted by calculating variance inflation factors (VIF) between each environmental parameter using the R package car (Fox & Weisberg, 2019) and GGally (Schloerke et al., 2024). Temperature (°C), salinity, depth (m), DO (µmol/kg), NO_3_ (µM) and FlECO-AFL (chlorophyll fluorescence, mg m^−3^) were selected for further analysis. A distance-based redundancy analysis (dbRDA) was performed, and ordinations were used to conduct ANOVA between environmental parameters and microbial ASVs using the vegan package. dbRDA plots were generated using ggplot2. Bray–Curtis distances were calculated from the fourth-root-transformed ASV relative abundance matrices. Five parameters were used for this analysis (temperature, salinity, DO, NO_3_, and FlECO-AFL) and only samples with complete environmental data were analyzed. The significance of the overall model, the marginal contribution of each variable, and each constrained axis were tested by permutational ANOVA with 999 permutations and the constrained variance was reported as the adjusted R². Ordinations were visualized with ggplot2, with the environmental variables displayed using stats::biplot.

## 3 Results

### 3.1 Hierarchical Clustering

The 78 samples were assigned to four sample clusters for this analysis (Figure 2). Out of 78 samples, 75 contained data for all nine parameters (temperature, salinity, DO, NO_3_, PO_4_, Sil, NH_4_, FlECO-AFL, and depth). PC1–PC4 captured 92.1 % of the variance. A top-down tree walk was performed to test for environmental differences among subclusters, but no internal node remained significant after Benjamini–Hochberg correction. We retained four clusters for further investigation defined by Bray–Curtis distance and Ward’s method clustering (k = 4), with Cluster FL1 corresponding to the 0.2 µm fraction and Clusters PA1–PA3 to large-sized fractions (3 and 20 µm). In addition, 221 filtered ASVs were grouped into six assemblages: Assemblage 1 (black; # of ASVs = 29), Assemblage 2 (pink; # of ASVs = 33), Assemblage 3 (yellow; # of ASVs = 30), Assemblage 4 (light blue; # of ASVs = 74), Assemblage 5 (dark blue; # of ASVs = 38), and Assemblage 6 (light red; # of ASVs = 17).

**Figure 2.**
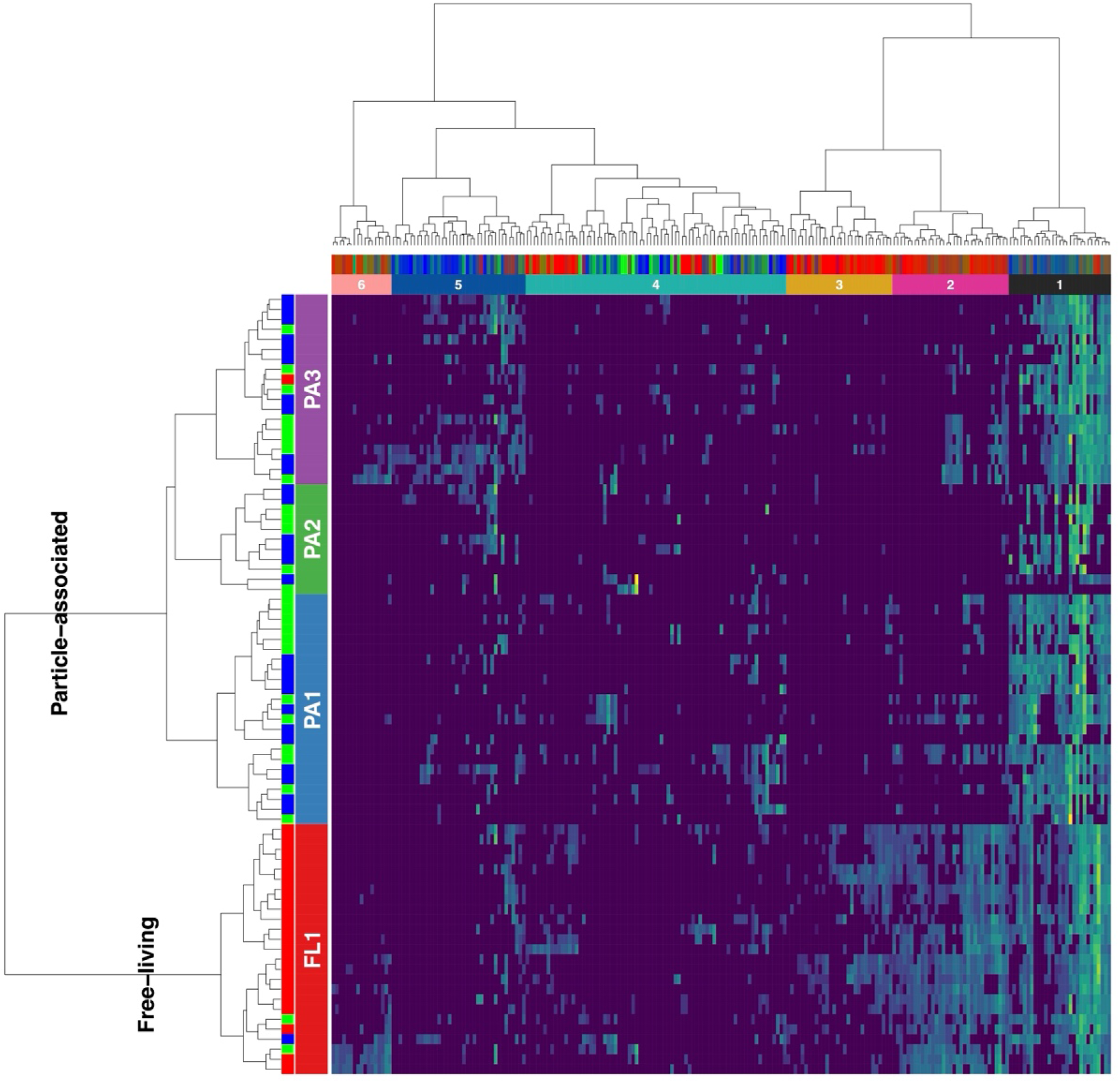
Heatmap of ASVs in the Process dataset. Hierarchical clustering was conducted based on relative abundance of 221 ASVs (columns), present in at least 5,000 quality-filtered reads across 78 seawater samples (rows), based on fourth-root–transformed relative abundances (viridis color scale; dark = low, yellow = high). Samples and ASVs were both clustered by Bray–Curtis dissimilarity with Ward’s method (ward.D). The dendrograms are shown on the left (samples) and top (ASVs). Samples were separated into four clusters: one free-living cluster (FL1) and three particle-associated clusters (PA1–PA3), indicated by the colored bar immediately left of the heatmap and by the labels within it. The first branch of the dendrogram (k = 2) separated the FL samples from the PA samples, as annotated on the tree. The second, thin row bar encodes the filter size of each sample (0.2 µm = red, 3 µm = green, 20 µm = blue). ASVs are colored by an RGB blend of their distribution across the three size fractions (red = 0.2 µm, green = 3 µm, blue = 20 µm), and are further grouped into six assemblages (numbered 1–6).

The relative abundances of the most abundant ASVs revealed differentiation in community composition across the four clusters (Figure 3), though ASVs from the three major bacterial classes were abundant throughout each of the four clusters. Alphaproteobacteria ASVs classified as SAR11 clade Ia are known to be free-living and were most abundant in Cluster FL1. Bacteroidetes ASVs classified as *Ulvibacter* are known to be common in algal phycospheres and were most abundant in particle-associated Clusters PA1–3. Gammaproteobacteria ASVs classified as uncultured Nitrincolaceae, SUP05, and SAR86 are known to be free-living and were most abundant in Cluster FL1. ASVs classified as *Psychrobacter* are common in sea ice and were unusually abundant in Cluster PA2.

**Figure 3.**
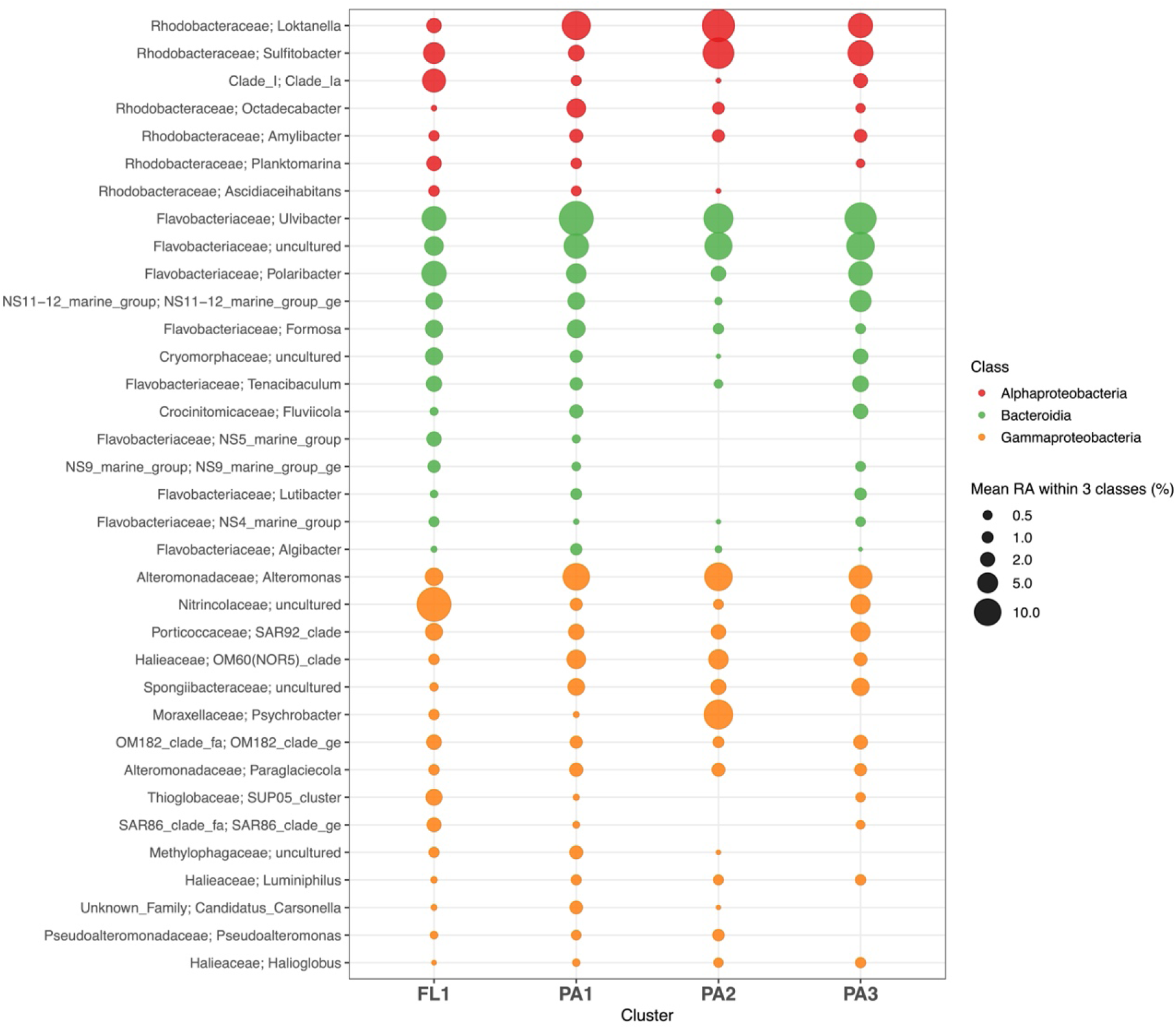
Bubble Plot of Major Genera Within Three Bloom-Responsive Bacterial Classes (Bacteroidia, Gammaproteobacteria, Alphaproteobacteria) Across the Four Clusters. Rows correspond to genera, columns to sample clusters. Bubble size represents the mean relative abundance (%) of the genus in each cluster in each class, and color represents class. Displayed genera satisfy two criteria: occurrence in > 10 % of samples (n = 78) and cumulative within-class relative abundance *≤* 95 %.

The three PA clusters shared a high relative abundance of ASVs belonging to three major bacterial classes. Cluster PA1 was dominated by *Ulvibacter*, *Loktanella*, and *Alteromonas*, with Bacteroidia being the most abundant class (42.4 %). Cluster PA2 showed the most evenly elevated abundances and contained ASVs not prominent elsewhere: *Psychrobacter*, *Sulfitobacter*, and *Alteromonas*. The abundance of *Psychrobacter* can be attributed to AnW and MWW which influenced this cluster. Also, this cluster lacked some Bacteroidia ASVs such as *Fluviicola*, NS5 and 9 marine groups, and *Lutibacter*, which were abundant in other clusters along with other Bacteroidia ASVs. Cluster PA3 was characterized by Bacteroidia ASVs including *Ulvibacter* and displayed the most diverse, including *Polaribacter*, uncultured Flavobacteriaceae, and the NS11-12 marine group. This cluster was AnW/cSW-influenced and comprised communities from the deepest samples, and Bacteroidia was the most abundant class (47.7 %), suggesting that it harbored taxa typical of particle-attached degraders of sinking diatom aggregates.

In the companion study of the survey stations (Shibue et al., 2026), the 11 survey clusters were assigned to different bloom stages. They were also significantly associated with water mass (Fisher’s exact test, p < 0.001), but the association was not one-to-one, as five clusters shared wSW as their primary water mass. Pre-bloom clusters tended to occur in wSW and post-bloom clusters in cSW, with one coastal cluster as an exception. On the other hand, in the process samples, there was no clear one-to-one correspondence between each water mass and bloom stage of the four clusters, as the clusters were primarily associated with the same water mass (AnW). However, each cluster differed in their bloom stages and the dominant bacterial class, driven by differences in size fractions.

The first split in the clustering dendrogram (Figure 2) discriminated free-living (FL; Cluster FL1; 0.2 µm filter fraction) and particle-associated (PA; Clusters PA1–3; 3 µm and 20 µm fractions). We did not use pore size to build the clusters. The samples separated this way based on their ASV composition alone. The partition of ASVs across the 0.2 µm, 3 µm, and 20 µm filter fractions (Figure 4) demonstrated size-selectivity of some ASVs. In particular, ASVs in Cluster FL1 were highly concentrated in the 0.2 µm fraction (most had > 60 % normalized mean relative abundance), while ASVs in Clusters PA1–3 overlapped substantially with each other and typically had < 40 % normalized mean relative abundance in the 0.2 µm fraction. This supports the result of hierarchical analysis (Figure 2) that separated process samples into FL (Cluster FL1) and PA (Clusters PA1–3) fractions and shows that the three PA clusters share particle-associated microbial community structures.

**Figure 4.**
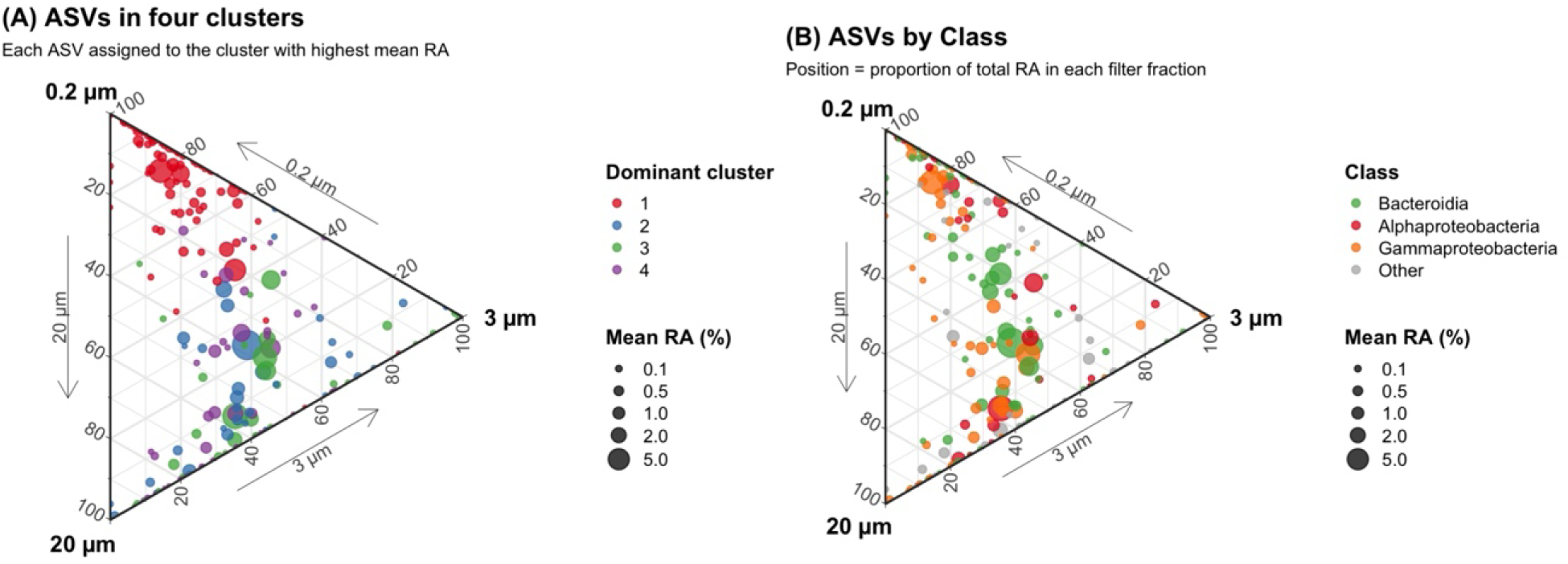
Ternary Diagrams of ASVs Across the 0.2, 3, and 20 µm Filter Fractions in the Process Dataset. Each point represents one of 221 ASVs that passed the abundance filter; its position on the ternary indicates the normalized mean relative abundance distributed among the 0.2 µm (free-living, top vertex), 3 µm (lower-right vertex), and 20 µm (bottom vertex) fractions, while point size encodes the mean within-sample relative abundance of the ASV across the 78 samples. (A) ASVs colored by the four sample clusters, defined as the cluster in which the ASV reached its highest mean relative abundance; (B) ASVs colored by taxonomic class: Bacteroidia (green), Alphaproteobacteria (red), and Gammaproteobacteria (orange); all other classes are pooled as “Other” (gray).

The three major bacterial classes were present in both FL and PA fractions (Figure 4; Table 2). However, the most abundant ASVs were rarely found to be specific to one size fraction and were typically intermediate between 2 or 3 size fractions. Gammaproteobacteria had higher mean abundance (40.4 %) in the 0.2 µm fraction than other fractions (29.1 % in 3 µm and 30.5 % in 20 µm), making them the most free-living-biased class of the three. Bacteroidia showed balanced portions of their abundance in each filter fraction, but the portion was relatively higher in 3 µm (36.0 %) than in other filter fractions (32.0 % in 0.2 µm and 20 µm). Alphaproteobacteria also showed balanced portions in each filter, but their abundance was somewhat higher in large-sized filter fractions than in 0.2 µm filter fraction (35.1 % in 20 µm, 34.3 % in 3 µm, and 30.6 % in 0.2 µm), driven by abundant Rhodobacteraceae members such as *Sulfitobacter* and *Loktanella*, known as bloom-responsive Alphaproteobacteria (Bergen et al., 2016; Sidhu et al., 2023). All three classes were present in every fraction, but each leaned toward a different one. Niche partitioning at the class level thus appears to come from abundance shifts, not from being absent from any fractions.

**Table 2.**
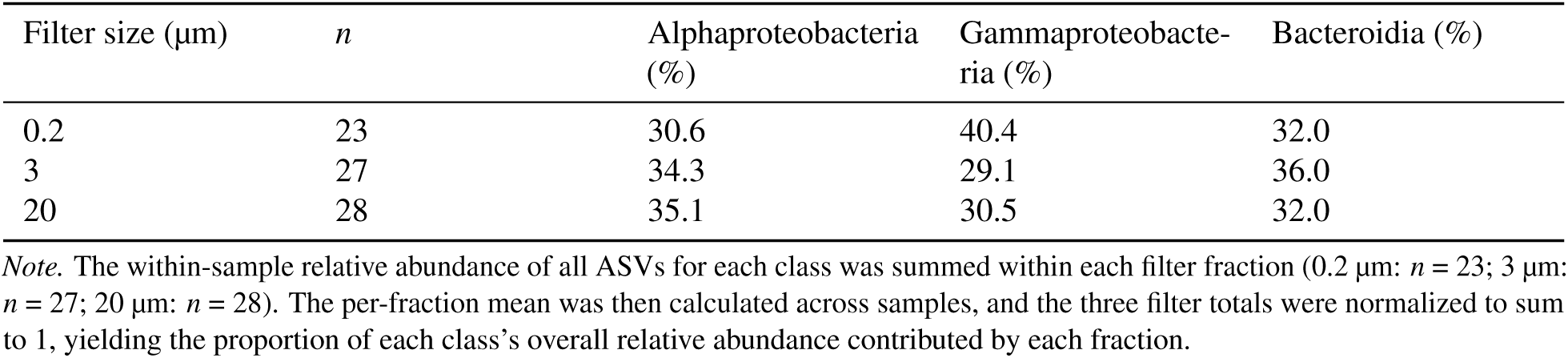
The Ratio of Three Bacterial Classes in Each Size Fraction.

ASVs in six assemblages were also investigated. As in the survey dataset, Assemblage 1 ASVs showed a cosmopolitan-like distribution, and their mean relative abundance was 74.5 % (Figure 5). However, compared to the survey dataset, six assemblages in the process dataset showed weaker biogeographical patterns as the number of samples is smaller than the survey dataset and the four sample clusters were separated based on FL and PA lifestyles, not the water mass differences.

**Figure 5.**
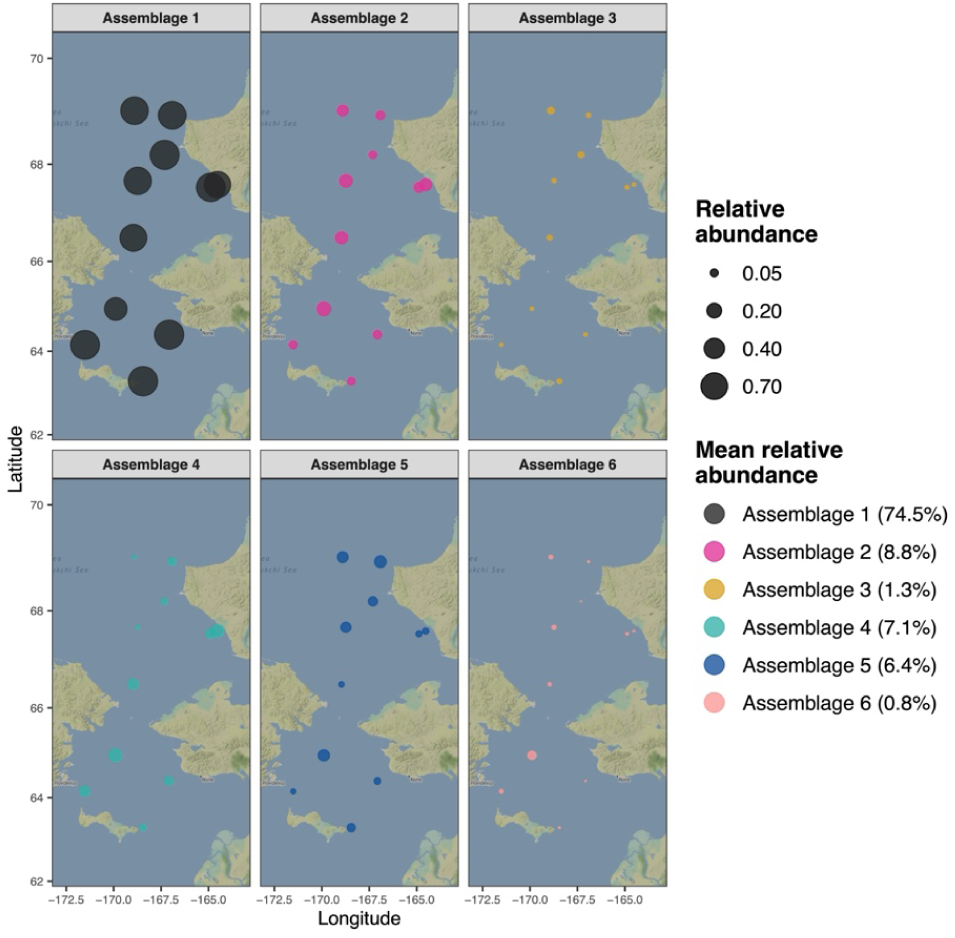
Sampling Sites of ASVs in Six Assemblages in the Process dataset. Each Assemblage is assigned to different colors. The dots correspond to mean relative abundance of ASVs in six assemblages. Mean relative abundance of ASVs in each Assemblage is shown as legend.

However, the bar plots showed different preferences toward different-sized filters (Figure 6). Assemblages 5 and 6 showed high abundance of ASVs (*≈* 50 %) among all size fractions. On the other hand, > 90 % of ASVs in Assemblages 2 and 3 were collected with 0.2 µm filter, suggesting that those assemblages were composed of FL communities. In contrast, Assemblage 4 was a PA community as ASVs were largely collected with 3 and 20 µm filters.

**Figure 6.**
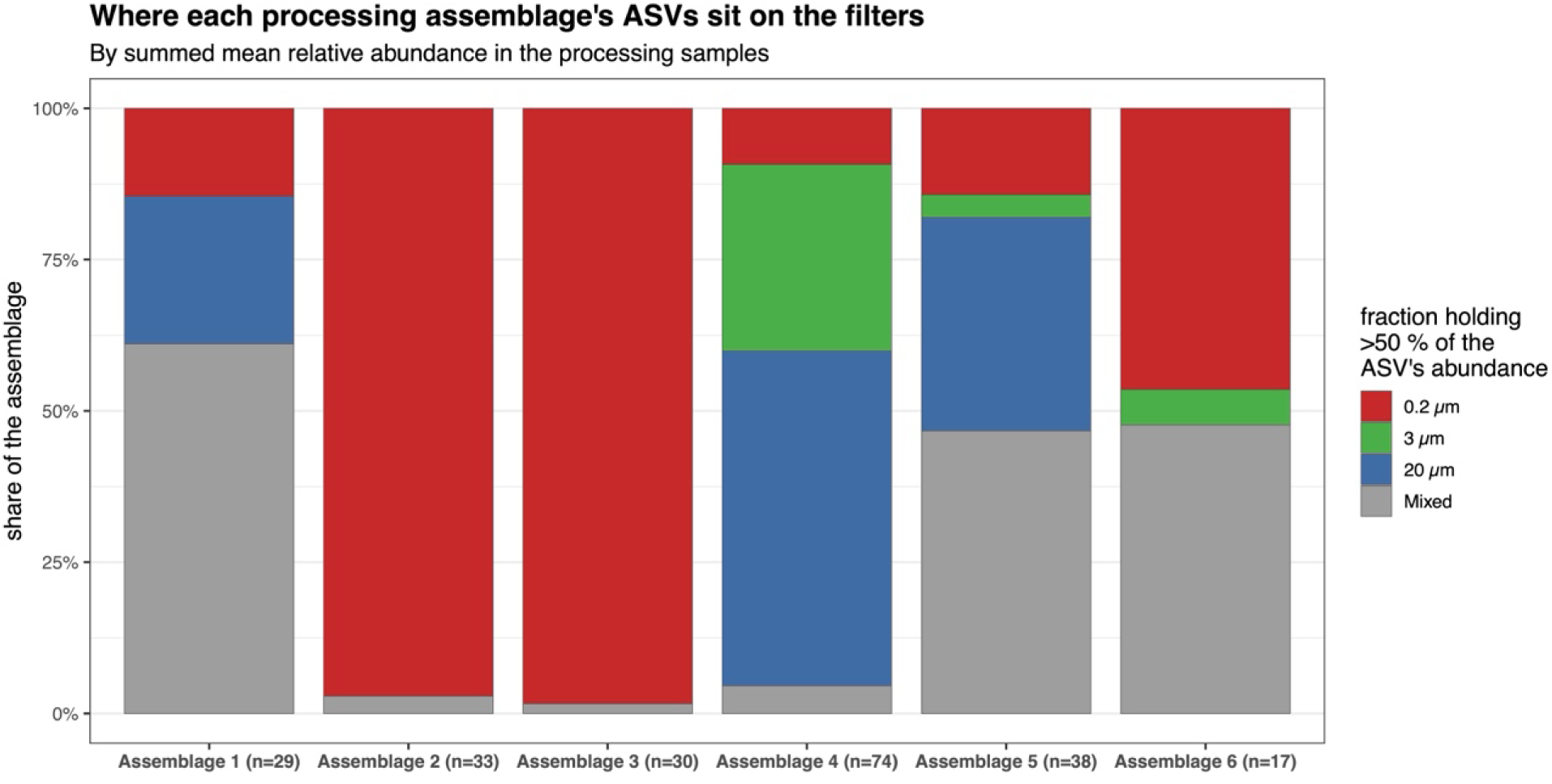
Relative Abundance of Six Assemblages ASVs With Each Filter in the Process Dataset. ASVs whose relative abundance was greater than 50 % were assigned to each filter size. Each filter is represented as different colors (red: 0.2 µm, green: 3 µm, blue: 20 µm). Gray color indicates ASVs whose abundance is spread across the size fractions, with no single filter holding more than 50 % of the ASV’s abundance.

Overall PERMANOVA confirmed that the four clusters differed strongly in composition (F = 15.08, R² = 0.379, p = 0.001), with the cluster factor explaining about 38 % of the total variation. However, PERMDISP was also significant (F = 7.46, p < 0.001), indicating that the four clusters did not have homogeneous multivariate dispersions. Pairwise PERMANOVA remained highly significant in every pair (BH-adjusted p = 0.001) with substantial effect sizes (R² = 0.19–0.39). The largest separations involved Cluster FL1 vs. the three PA clusters (R² = 0.33–0.39 for FL1 vs PA1–PA3), while the three PA clusters differed less from each other (R² = 0.19–0.22 for the PA1 vs PA2, PA1 vs PA3, and PA2 vs PA3 pairs). On the other hand, pairwise PERMDISP showed that the dispersion difference came from Cluster PA2. Cluster PA2 had significantly greater dispersion than Cluster FL1 (Tukey’s HSD, p < 0.001), Cluster PA1 (Tukey’s HSD, p = 0.011), and Cluster PA3 (Tukey’s HSD, p = 0.010), whereas pairwise dispersion among Clusters FL1, PA1, and PA3 was statistically homogeneous. In summary, the four clusters represent genuinely distinct communities, based on three pieces of evidence: (1) the overall PERMANOVA has a large R^2^; (2) every pairwise comparison is significant; (3) Clusters FL1, PA1, and PA3 have similar within-cluster dispersion, so dispersion itself is not driving the result. Cluster PA2 has a distinct mean position from the other clusters but is internally heterogeneous, suggesting that this cluster forms a more diverse set of communities than the other three clusters.

### 3.2 Geographic and Hydrographic Distribution of Sample Clusters

There were not strong gradients in vertical or horizontal spatial distributions of the sample clusters (Figure 7). Cluster FL1 (n = 25) was the most geographically widespread, present at all 11 sampling stations (4–12 % per station), indicating that microbial communities in the 0.2 µm fraction possibly occur throughout the study area regardless of location. The three PA clusters showed small station-level differences that nevertheless spanned both seas. Cluster PA1 (n = 23) was concentrated at stations in IL and CBW Transects (22 % in IL1.1, and 17 % in CBW5) and was absent from DBO stations (DBO2.4 and DBO3.8A). Cluster PA2 (n = 11) was concentrated along DBO Transect (18 % each at CBE9, DBO2.4, DBO3.3) and was absent from stations in CBW, CL1, CNL, and IL Transects. Cluster PA3 (n = 19) was concentrated in the northern Bering and the northern Chukchi Seas (16 % each at CBE1, CL1, CL3, and DBO2.4) and absent from stations in CBW, CNL, and IL Transects. More than 60 % of samples in Clusters FL1, PA1, and PA3 were collected from the Chukchi Sea (lat *≥* 65.7°N), while samples in Cluster PA2 were evenly collected from both Bering and Chukchi Seas (50 % each), implying that high within-cluster dispersion in Cluster PA2 can be attributed to this result.

**Figure 7.**
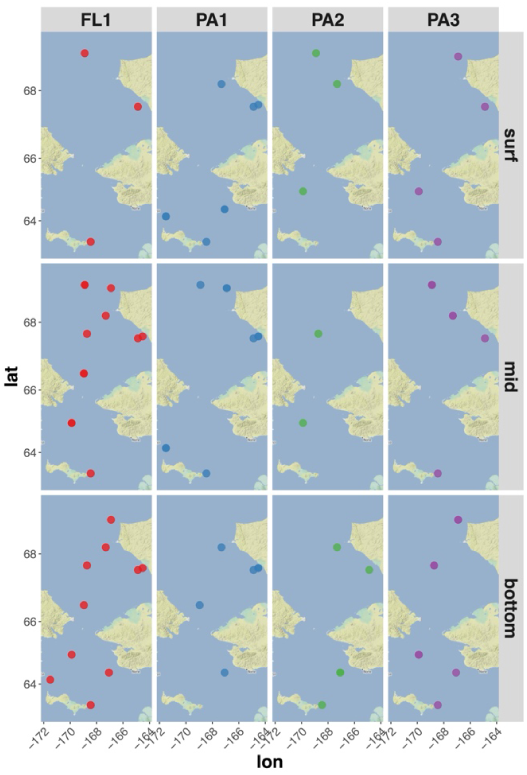
Maps of Sampling Stations for Each of the Four Sample Clusters. Each dot is colored by the four clusters: Cluster FL1 (red; n = 25), Cluster PA1 (blue; n = 23), Cluster PA2 (green; n = 11), and Cluster PA3 (purple; n = 19). One PA2 sample came from a shipboard incubation without a station location and is not mapped. Each map is separated by depth (surface, middle, and bottom).

No clear differences in water mass characterizations were observed across the four sample clusters, with each cluster found in multiple water masses (Figure 8; Table 3). Clusters FL1 (FL mid-bottom cluster) and PA3 (PA mid-bottom cluster) were both dominated by AnW (48 % and 47 %, respectively), with warm Shelf Water (wSW) as the secondary water mass in Cluster FL1 and cool Shelf Water (cSW) in Cluster PA3. Cluster PA2 (PA deep cluster) was also AnW-dominated (40 %) but had the highest proportional MWW content of any cluster (20 %). On the other hand, Cluster PA1 (surface PA cluster) was the only cluster whose primary water mass was wSW (48 %), reflecting its warmer and lower-salinity character, with Modified Winter Water (MWW) emerging as a secondary water mass (13 %).

**Figure 8.**
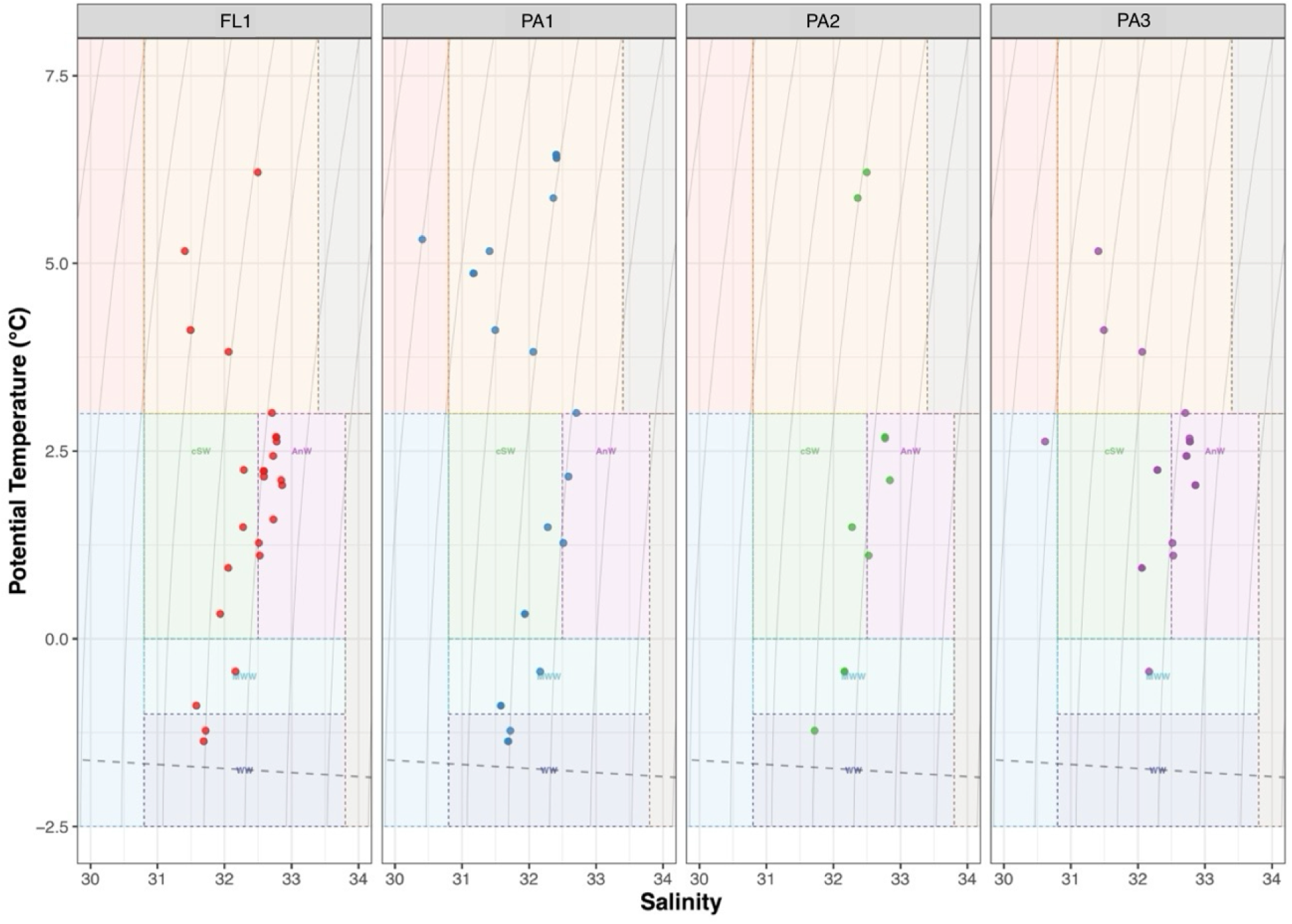
T-S Diagrams With Water Mass Boundaries for Each of the Four Clusters. Dashed rectangles indicate the T-S ranges of each water mass: warm Coastal Water (wCW), cool Coastal Water (cCW), Ice Melt Water (IMW), warm Shelf Water (wSW), cool Shelf Water (cSW), Anadyr Water (AnW), Modified Winter Water (MWW), Winter Water (WW), Atlantic Water (AtlW), Bering Basin Water (BBW). Gray contours represent isopycnals (lines of equal density, *σ_θ_*, at 0.5 kg/m³ intervals). The dashed black line indicates the freezing point.

**Table 3.** The Proportion of Samples in the Four Clusters, Classified Into Each Water Mass.

| Cluster | Primary water mass | Secondary water mass |
|---|---|---|
| FL1 | AnW (48 %) | wSW (20 %) |
| PA1 | wSW (47.8 %) | MWW (13 %) |
| PA2 | AnW (40 %) | MWW (20 %) |
| PA3 | AnW (47.4 %) | cSW (21.1 %) |
| Fisher’s exact test (Monte Carlo) |  |  |
| Replicates | *p* | Significance |
| B = 9,999 | = .126 | ns |

*Note.* The primary water mass was defined as the water mass with the highest proportion of samples, and the secondary water mass as the one with the second highest proportion. Water-mass abbreviations are defined in Table 1. *ns* = not significant.

A total of 78 samples were collected from the Bering Sea (n = 29) and the Chukchi Sea (n = 48) from different depths: surface (n = 9), middle (n = 8), and bottom (n = 12) in the Bering Sea; surface (n = 9), middle (n = 22), and bottom (n = 17) in the Chukchi Sea. The FL and PA communities did not significantly differ with depth. The PA community accounted for most of the microbial community composition in surface waters (mean: 73.2 %) and declined at mid and bottom depths (mean: 59.7 % and 61.7 %, respectively), while the FL community increased from 26.8 % at the surface to 40.3 % and 38.3 % at mid and bottom depths. However, a Kruskal–Wallis test revealed that this proportional difference of bacterial ASVs in each depth was not statistically significant (*χ*² = 1.95, df = 2, p = 0.38). This is consistent with the result from the survey samples and can be attributed to the shallowness and the weak stratification in the Bering and Chukchi Seas during June 2017 (Figure 14 in Shibue et al., 2026).

Figure 9 is the map showing the depth profile of ASV richness in the FL and PA clusters. Samples from each cluster were mainly collected from the Chukchi Sea, except Cluster PA2 whose samples were evenly collected from both Bering and Chukchi Seas (5 samples each). In most clusters, the station with the highest ASV richness shifted with depth. In Cluster FL1, ASV richness peaked at CL3 (74 observed ASVs) at the surface, at CL1 (90) at mid-depth, and at IL2 (74) at the bottom. In Cluster PA1, richness peaked at CBE9 (65 observed ASVs) at the surface, at CBW5 (64) at mid-depth, and at IL1.1 (67) at the bottom, with little change in the peak value. In Cluster PA2, the maximum remained among the DBO stations (DBO2.4, DBO3.8A, and DBO3.3) at all depths but stayed at consistently low richness (32–36 observed ASVs). In Cluster PA3, richness peaked at IL2 (49) and CL3 (50) at the surface and mid-depth and reached its highest value at DBO3.8A (74) at the bottom.

**Figure 9.**
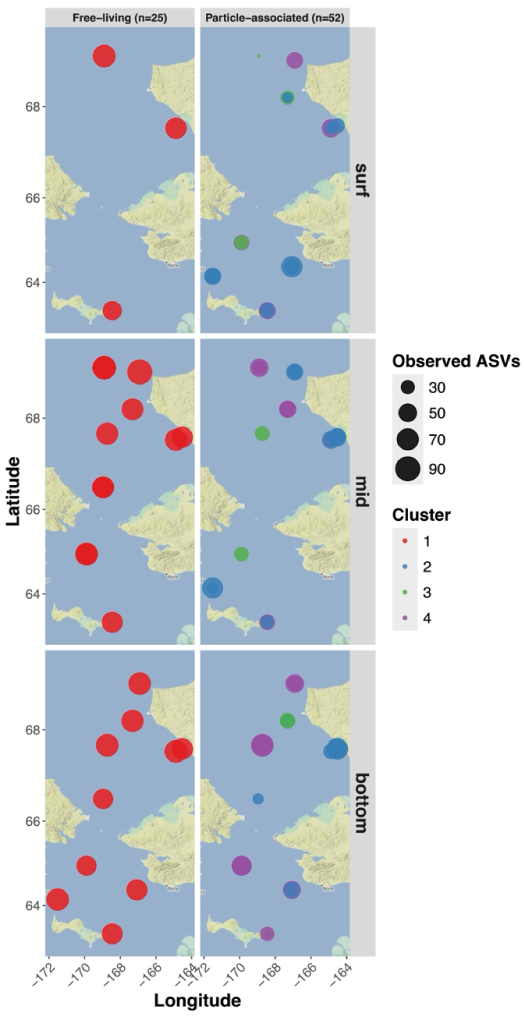
ASV Richness in Free-Living and Particle-Associated Communities, Shown by Depth (Surface, Middle, and Bottom). Each point represents a sampling location.

However, along with the depth profiles of the FL and PA communities, Kruskal–Wallis tests were conducted to assess whether ASV richness (observed ASVs) differed in each depth and showed no significant depth effect on richness for any cluster after Bonferroni correction (*α* = 0.0125): Cluster FL1 (*χ*² = 1.87, df = 2, p = 0.392), Cluster PA1 (*χ*² = 0.42, df = 2, p = 0.810), Cluster PA2 (*χ*² = 2.08, df = 2, p = 0.354), and Cluster PA3 (*χ*² = 4.90, df = 2, p = 0.086). Although Cluster PA3 tended to show higher richness at the bottom (median 53.5 observed ASVs) than at the surface and mid depths (40.5 and 36.0, respectively), this trend did not remain significant after Bonferroni correction for multiple comparisons. Across all samples, richness likewise did not differ significantly among depths (*χ*² = 3.07, df = 2, p = 0.215). These results indicate that ASV richness was similar across depths in this region during the sampling period. However, depth should not be regarded as unimportant. Salinity and nutrient concentrations still changed with depth, and it is through these gradients that depth acts here.

In survey samples, Bacteroidia showed high abundance at DBO3 stations and other transects nearby (Table 3 in Shibue et al., 2026). This can be explained by the active diatom bloom caused by *Chaetoceros* and *Thalassiosira* at DBO3 stations during June 2017. On the other hand, in process samples, two sample stations were in DBO3 Transect (DBO3.3 and 3.8A stations), and each cluster evenly had samples from both stations (Table 4). Samples from CBW Transect and other transects near DBO3 stations showed high abundance of Bacteroidia, while DBO3 stations showed relatively lower abundance of this class compared to those transects. This contrasts with the survey dataset (> 35 %) at DBO3 stations and nearby transects but can be explained by the difference of sample sizes and locations; 30 samples from 8 DBO3 stations were collected in survey samples, while 13 samples from 2 DBO3 stations were collected in process samples. However, though Bacteroidia were proportionally lower at DBO3 than at other process transects, it still accounted for a substantial fraction of the community (32.6 %), the second most abundant class after Gammaproteobacteria, and only marginally below the *≥* 35 % level observed in survey samples. Also, at the family–genus level, ASVs belonging to Bacteroidia, such as *Ulvibacter*, uncultured Flavobacteria, *Polaribacter*, and NS11-12 marine group, accounted for the majority of the highly abundant ASVs, whereas the high abundance of Gammaproteobacteria was driven largely by a single genus, *Alteromonas* (16.7 %). Therefore, the two datasets differed at DBO3 in the relative abundance of these classes, not in which taxa dominated. In contrast to the survey, where the divisions were based on water mass and hierarchical clustering, the four clusters in process samples were separated by filter size fractions. Nevertheless, the high relative abundance of the major bacterial classes was likewise confirmed, suggesting that the consistent high abundance of those major bacterial classes was recovered during the ASGARD 2017 cruise regardless of how the communities were partitioned.

**Table 4.**
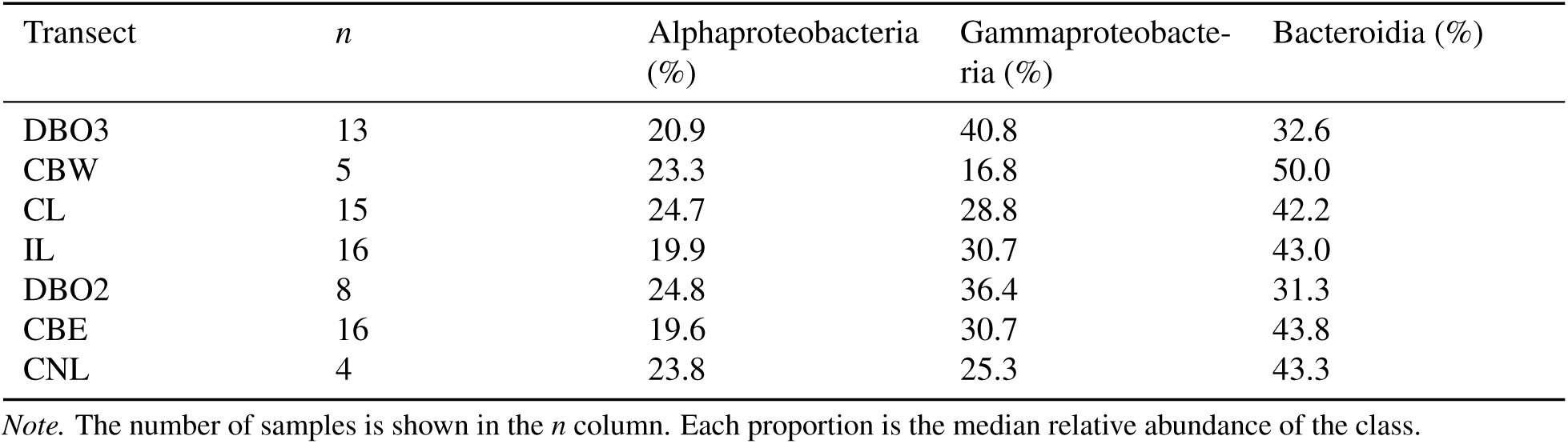
Composition of the Dominant Three Bacterial Classes (Alphaproteobacteria, Gammaproteobacteria, and Bacteroidia) Across the Seven Transects.

| Transect | *n* | Alphaproteobacteria (%) | Gammaproteobacte-ria (%) | Bacteroidia (%) |
|---|---|---|---|---|
| DBO3 | 13 | 20.9 | 40.8 | 32.6 |
| CBW | 5 | 23.3 | 16.8 | 50.0 |
| CL | 15 | 24.7 | 28.8 | 42.2 |
| IL | 16 | 19.9 | 30.7 | 43.0 |
| DBO2 | 8 | 24.8 | 36.4 | 31.3 |
| CBE | 16 | 19.6 | 30.7 | 43.8 |
| CNL | 4 | 23.8 | 25.3 | 43.3 |

*Note.* The number of samples is shown in the *n* column. Each proportion is the median relative abundance of the class.

### 3.3 Physicochemical Characterization of Sample Clusters

No significant differences (p *≥* 0.05) in physicochemical characteristics were observed across clusters using six environmental variables (Figure 10). Cluster FL1 was characterized by an intermediate temperature (median: 2.16 °C), an intermediate sampling depth (median: 20.1 m), and the highest NO_3_ of any cluster (median: 2.60 µM). Among the three PA clusters, Cluster PA1 was the most distinct: it had the highest temperature (mean: 3.01 °C), the lowest salinity (mean: 31.9), the highest dissolved oxygen (mean: 352.4 µmol kg^−1^), and the shallowest sampling depth (mean: 15.9 m), all consistent with wSW-influenced post-bloom surface conditions. Cluster PA2 had cold (mean: 1.80 °C), high-salinity (mean: 32.4) deep water and showed the highest FlECO-AFL fluorescence (mean: 2.00 mg m^−^³). Cluster PA3 occupied the deepest samples (mean: 25.9 m) and had the lowest NO_3_ (mean: 0.60 µM) and one of the lowest FlECO-AFL values (mean: 0.86 mg m^−3^), suggestive of a post-bloom community.

**Figure 10.**
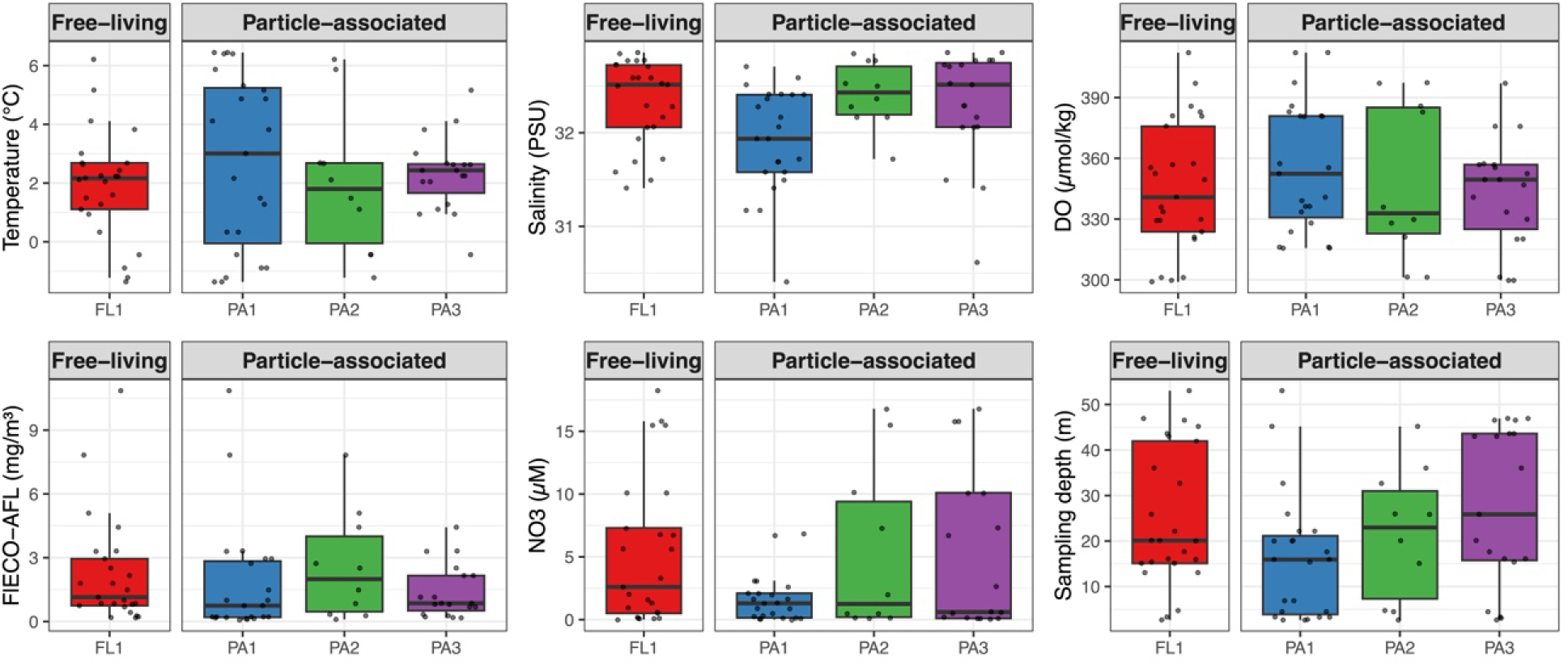
Boxplots of Environmental Variables for the Four Clusters, Grouped by Free-Living (FL) and Particle-Associated (PA) Clusters. Each panel shows one of six variables: temperature (°C), salinity (PSU), DO (µmol/kg), FlECO-AFL (chlorophyll fluorescence, mg m^−3^), NO_3_ (µM), and sampling depth (m). Individual samples were overlaid as jittered points. Boxes represent the interquartile range (IQR), with the horizontal line indicating the median. Whiskers extend to the most extreme values within 1.5 × IQR.

The bloom index (rescaled PC1) was not significantly different across the four clusters (p *≥* 0.05) (Figure 11). The median bloom index was highest in Cluster PA1 (0.58), suggestive of Cluster PA1 as a post-bloom community characterized by nutrient-scarce warmer shallower water. The three other clusters showed lower bloom indices (0.39 in Cluster FL1; 0.44 in Cluster PA2; 0.46 in Cluster PA3). Cluster FL1 had the lowest bloom index, consistent with a free-living community persisting in nutrient-replete, mid-depth waters. Clusters PA2 and PA3 occupied intermediate bloom index values; Cluster PA2 had the highest FlECO-AFL concentration, and Cluster PA3 had the lowest FlECO-AFL and NO_3_ concentrations, suggestive of during-bloom and post-bloom conditions, respectively.

**Figure 11.**
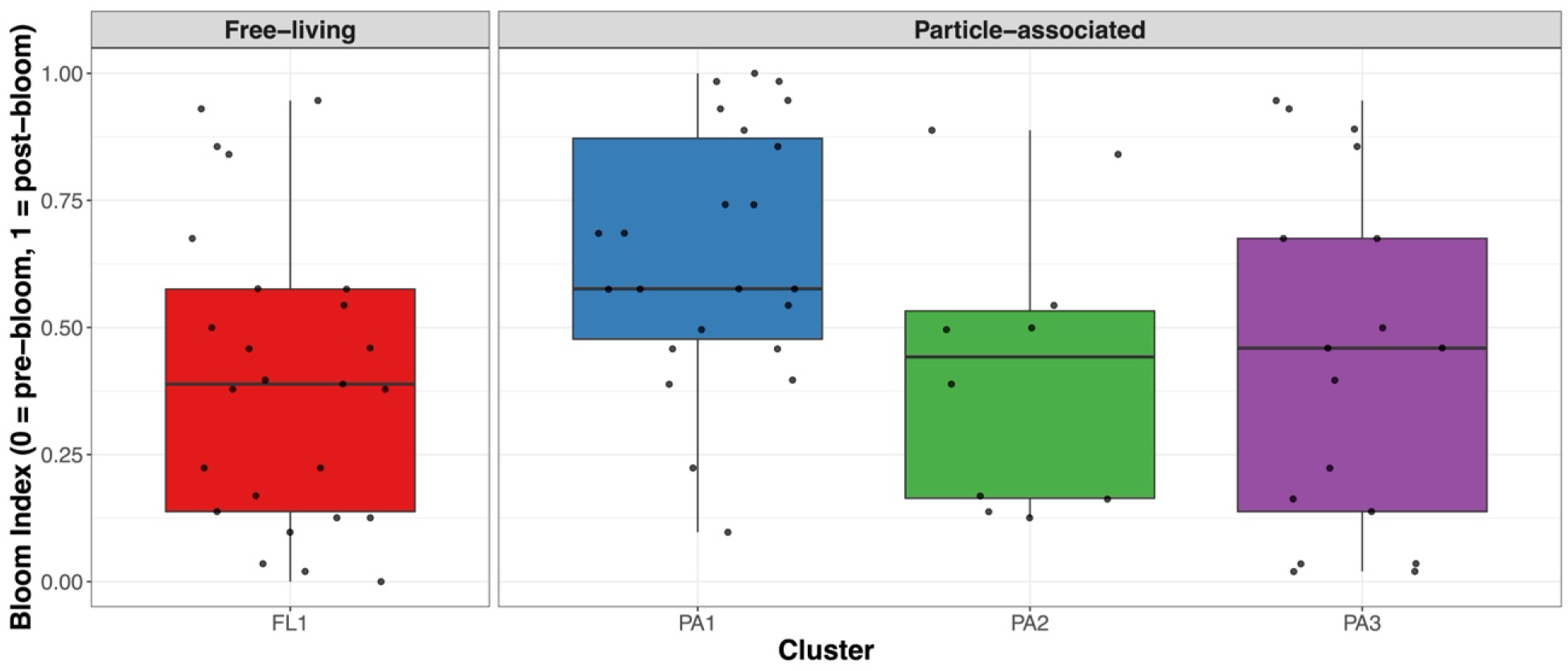
Bloom Index (Rescaled PC1) for the Four Sample Clusters, Grouped by Free-Living (FL) and Particle-Associated (PA) Clusters. The bloom index was derived from PC1 of a PCA on nine environmental variables, rescaled from 0 (pre-bloom, nutrient-replete) to 1 (post-bloom, nutrient-depleted). Boxes represent the interquartile range with the median shown as a horizontal line. Individual samples are overlaid as jittered points.

Ordination using dbRDA also failed to reveal strong relationships between cluster community structure and five selected environmental parameters (temperature, salinity, DO, NO_3_, and FlECO-AFL; Figure 12). The constrained ordination model was statistically tested by permutational ANOVA (999 permutations).

**Figure 12.**
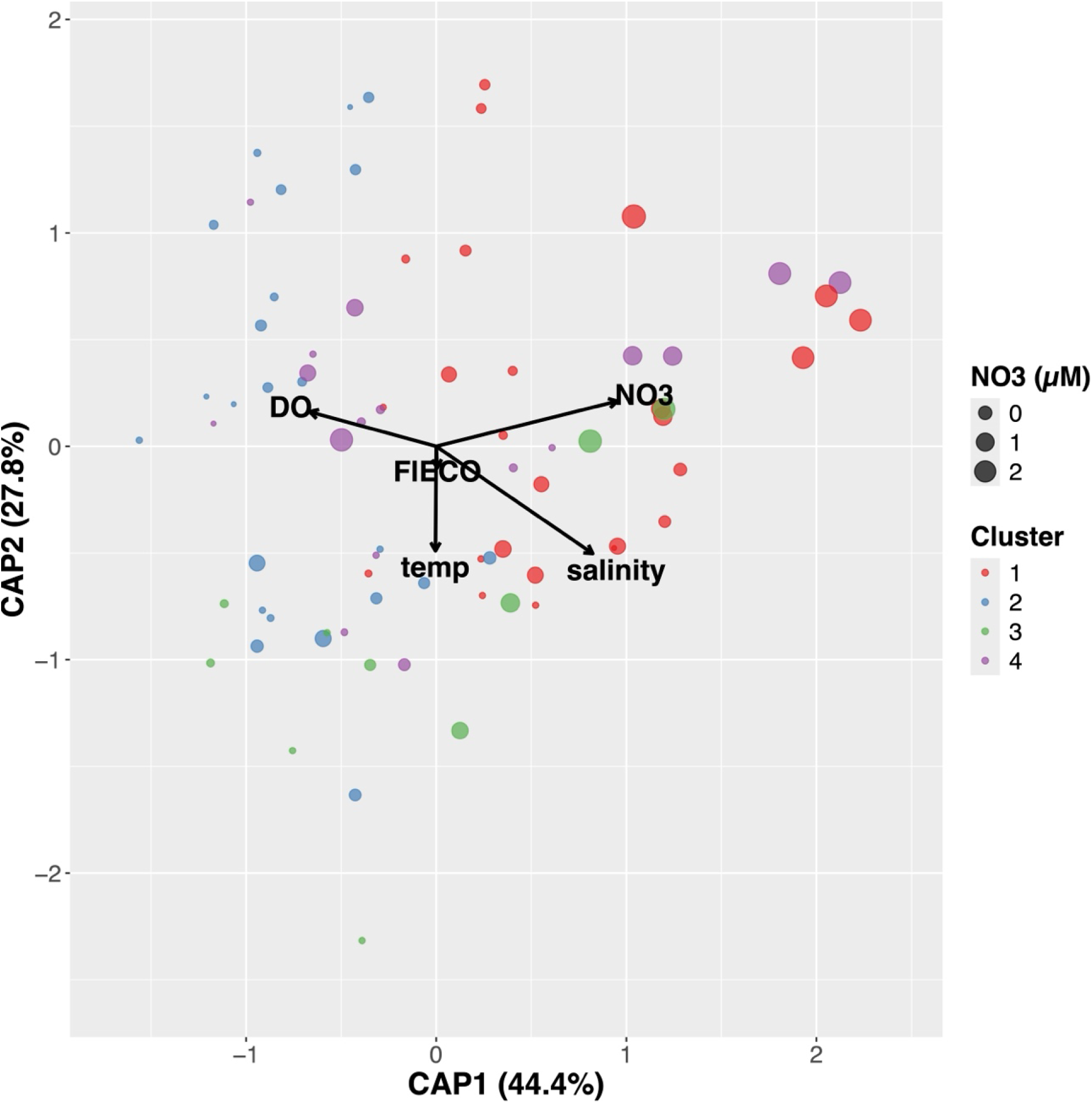
Distance-Based Redundancy Analysis (dbRDA) of Microbial Community Composition Constrained by Five Environmental Variables. Temperature, salinity, DO, NO_3_, and FlECO-AFL were used. Samples are plotted on the first two canonical axes (CAP1, 44.4 %; CAP2, 27.8 %), colored by the four process clusters, with dot size scaled to NO_3_ concentration (µM). Arrows indicate the direction and relative strength of each environmental variable’s contribution to the constrained ordination.

The overall dbRDA was highly significant (F = 2.66, p = 0.001), with the five environmental parameters collectively explaining only 16.2 % of variation in community composition (adjusted R² = 0.10). Among individual variables, only salinity had a significant marginal effect (F = 3.15, p = 0.004); FlECO-AFL fluorescence, temperature, dissolved oxygen, and NO_3_ were not significant. Of the five constrained axes, only CAP1 (F = 5.91, p = 0.002) and CAP2 (F = 3.76, p = 0.031) were significant, justifying the use of the first two axes for biplot interpretation. Together these results indicate that process-site community variation was structured primarily along a salinity axis, with a smaller contribution from FlECO-AFL. CAP1 (44.4 % of constrained variance) was positively associated with NO_3_ (+0.95) and high salinity (+0.82), and negatively with dissolved oxygen (−0.67), capturing a nutrient–salinity gradient: nutrient-rich, salty, oxygen-poor deep water on one side, and nutrient-poor, fresher, oxygen-rich surface water on the other. CAP2 (27.8 % of constrained variance) was driven mostly by temperature (−0.49) and salinity (−0.50), with a small positive contribution from DO (+0.16), capturing a temperature–salinity gradient, separating warmer, fresher shelf water from colder, saltier source water at depth.

On the CAP ordination, the four clusters separated along ecologically meaningful gradients. Cluster FL1 was placed at high CAP1, driven by high NO_3_ and salinity. Cluster PA1 sat at low CAP1, driven by low NO_3_ and salinity, and high dissolved oxygen, consistent with a post-bloom surface condition. Cluster PA2 was located at the lowest CAP2, where its centroid was pulled by relatively low DO and temperature contrast captured by CAP2. Cluster PA3 sat near the center of the plot, slightly offset toward Cluster FL1, reflecting a community that shares many features with the other deep-water clusters but with a less distinct environmental signature.

The equivalent PCA for the process samples (n = 75) recovered essentially the same structure (Figure 13). PC1 explained 48.6 % and PC2 24.8 % of the variance (73.4 % cumulative). As in the survey dataset, PC1 separated positive loadings for PO_4_ (0.45), NO_3_ (0.43), NH_4_ (0.37), silicate (0.35), salinity (0.34) and depth (0.32) from a negative loading for temperature (−0.22), and PC2 was again dominated by dissolved oxygen (0.59) versus temperature (−0.56). The process samples likewise formed a curved arch along PC1, reproducing the horseshoe/arch pattern of a single continuous gradient. Therefore, both datasets shared almost identical variance partitioning (PC1 *≈* 48 % in both datasets), loadings and arch shape. This indicates that the survey and process samples are governed by the same primary physicochemical gradient and the two datasets are directly comparable despite their different spatial coverage and sample numbers.

**Figure 13.**
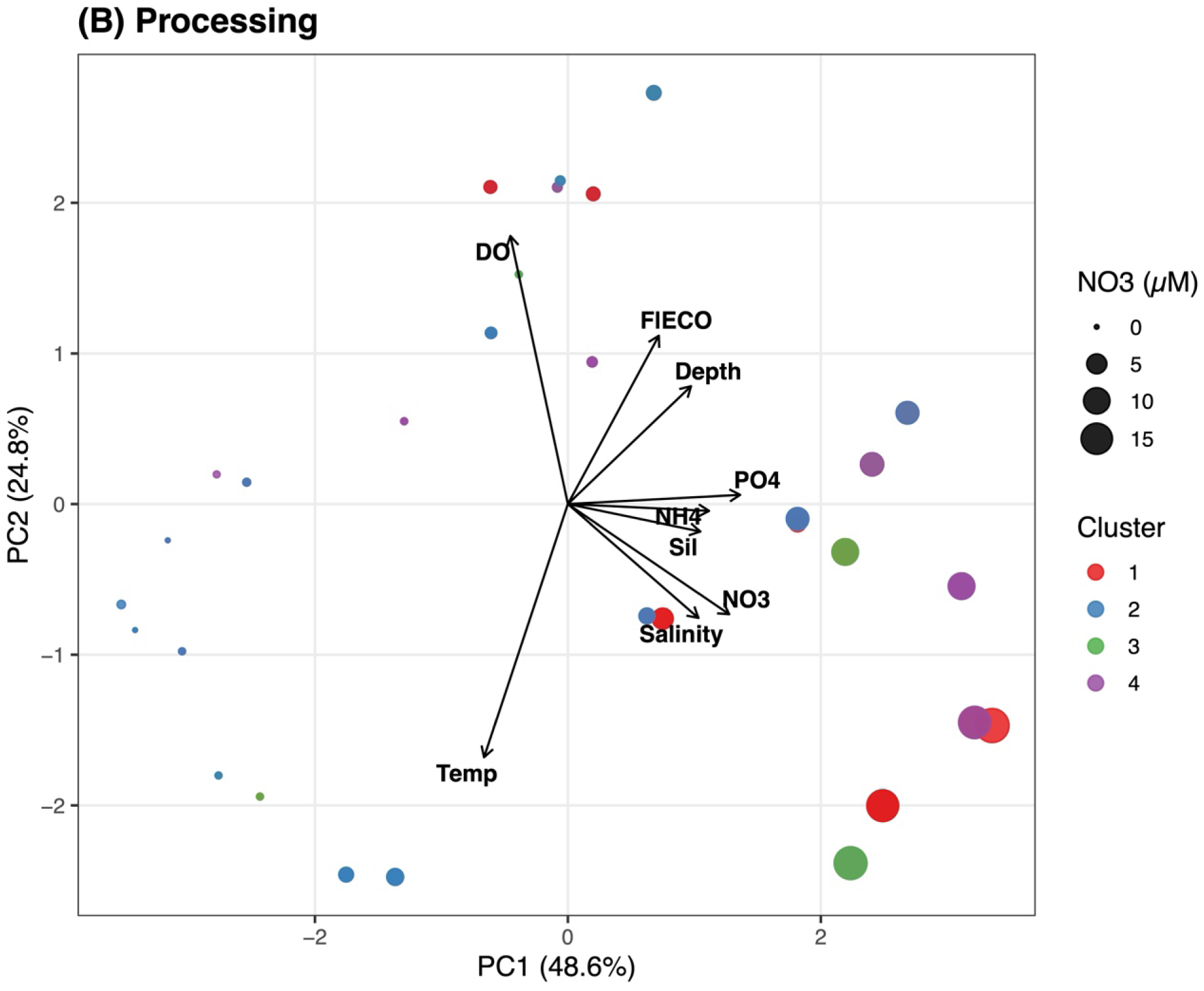
PCA Biplots of the Survey and Process Datasets. Principal component analysis (PCA) of the environmental data of the survey (A) and the process (B) data is shown. Nine variables (temperature, salinity, dissolved oxygen, NO_3_, PO_4_, silicate, NH_4_, fluorescence and depth) were used, with nutrients and fluorescence log-transformed and all variables standardized prior to PCA. Each point is a sample, positioned by its PC1 and PC2, colored by sample clusters, and sized by nitrate concentration (µM). Black arrows show the loadings of the environmental variables. Axes are labelled with the percentage of variance explained (survey: PC1 48.1 %, PC2 20.7 %, n = 174; process: PC1 48.6 %, PC2 24.8 %, n = 75). PCA was computed separately for each dataset, so scores are not directly comparable between them.

## 4 Discussion

### 4.1 Ecological Characterization

In our study, contrary to the survey dataset, six assemblages were characterized by different lifestyles (Figures 4 and 6). Also, ASVs belonging to Alphaproteobacteria, Gammaproteobacteria, and Bacteroidia in the process dataset accounted for 93.9 % of the total relative abundance, consistent with previous work in this region (Chen et al., 2008; Lee et al., 2019; Zeng et al., 2012; Zeng et al., 2013a), as well as with our survey dataset (94.7 %). However, the relative importance of these classes differed between datasets. Gammaproteobacteria was the most abundant class in the survey dataset, whereas Bacteroidia was the most abundant (40.2 %) in the process dataset. This is likely because the process dataset was size-fractionated by sequential filtration into free-living and particle-associated samples, and Bacteroidia are strongly particle-associated, attaching to and breaking down algal particles. Gammaproteobacteria, which are more common in the general community, were most abundant in the survey dataset. Bacteroidia was abundant across all size fractions, including the 0.2 µm free-living fraction (38.5 %), and not only in the 3 and 20 µm particle-associated fractions. Because Bacteroidetes are closely associated with algal blooms and with the degradation of high molecular weight compounds produced by phytoplankton (Bunse & Pinhassi, 2017; Needham & Fuhrman, 2016; Needham et al., 2018), the higher Bacteroidia abundance in the process dataset may reflect the bloom-influenced conditions of the process stations.

A pyrosequencing study in the Chukchi Sea, the Beaufort Sea, and Franklin Bay reported that Gammapro-teobacteria and Alphaproteobacteria together accounted for 22–30 % of all sequences, and that the two most abundant clusters at 97 % similarity were SAR11 and one belonging to Bacteroidetes (Kirchman et al., 2010). Using amplicon sequencing across the Arctic including the Chukchi Sea, and the Antarctic, Ghiglione et al. (2012) found that most OTUs in both polar oceans belonged to Gammaproteobacteria, Alphaproteobacteria, and Bacteroidia, along with ubiquitous, metabolically versatile OTUs such as SAR11 and SAR324 (Malfertheiner et al., 2022); *Polaribacter*, a representative bloom-associated bacterium, was among the Arctic-specific OTUs. Pedrós-Alió et al. (2015) further reported that temperate communities were dominated by Alphaproteobacteria (approximately 50 %), whereas Arctic communities held nearly equal proportions of Alphaproteobacteria and Gammaproteobacteria (about 20 % each), with Bacteroidetes exceeding the global average (18 % vs. 4 %). The reasons for the high abundance of Bacteroidetes in polar oceans remain unclear, although the quantity and quality of available substrates are thought to be involved (Kirchman et al., 2007).

Metagenomic analyses have similarly documented the dominance of Alphaproteobacteria, Gammapro-teobacteria, and Bacteroidetes in the Arctic Ocean. Cao et al. (2020), examining metagenomes from the poles to the temperate region, showed that 90 phylum-level taxa classified from 24,504 OTUs mostly belonged to Alphaproteobacteria, Gammaproteobacteria, and Bacteroidetes, which were highly abundant in both surface and deep-water Arctic samples. Royo-Llonch et al. (2021) analyzed 530 MAGs reconstructed from the Arctic Ocean and found that more than half belonged to Gammaproteobacteria (22.8 %), Bacteroidota (20.9 %), and Alphaproteobacteria (11.5 %), while the 12 most abundant MAGs represented unknown species within those groups and other phyla such as Actinobacteria and SAR324.

The association between bacteria and algal blooms has been studied from temperate regions to the Arctic Ocean. In the North Sea, the SAR11 clade was abundant in pre-bloom communities, Bacteroidetes such as *Formosa*, *Polaribacter*, and *Ulvibacter* during the bloom, and Gammaproteobacteria such as the SAR92 clade post-bloom (Sintes et al., 2013; Teeling et al., 2012; Teeling et al., 2016). Although those results were from temperate regions, these patterns are consistent with our results. In Franklin Bay, Bacteroidetes and Al-phaproteobacterial OTU *Roseobacter*, which is a major marine bacterium contributing to carbon cycling and aerobic anoxygenic photosynthesis (Brinkhoff et al., 2008), peaked at the chlorophyll a maximum, whereas Gammaproteobacteria became abundant afterward (Alonso-Sáez et al., 2008). Through the northern Barents Sea into the area east of Svalbard, Gammaproteobacteria ASVs (e.g., SAR92, OM60, and Nitrincolaceae) and Bacteroidetes ASVs (e.g., *Formosa*, *Polaribacter*, and the NS9 marine group) became abundant during the spring–summer sea-ice algal bloom, while SAR11 and some nitrifiers dominated winter samples (Thiele et al., 2023). A time-series study from the Dease Strait assigned 80 % of reads to Alphaproteobacteria (33.2 %), Gammaproteobacteria (28.7 %), and Bacteroidia (21.6 %) (Redondo-Río et al., 2024), and Alphaproteobacteria dominated the pre-bloom community, Bacteroidetes rose during the bloom, and Gammaproteobacteria increased over the bloom but declined once Bacteroidetes rose exponentially.

Several Arctic studies have described the taxonomic composition of FL and PA bacterial communities. In surface waters along the Mackenzie River–Beaufort Sea transects (Canadian Arctic), Ortega-Retuerta et al. (2013) found that the offshore PA fraction was dominated by Bacteroidetes and Gammaproteobacteria, whereas the FL fraction was dominated by Alphaproteobacteria. In the river-influenced Laptev Sea, Kellogg and Deming (2009) separated FL (0.22–1 µm), suspended-particle (1–60 µm), and aggregate (>60 µm) fractions and reported bacterial libraries dominated overall by Gammaproteobacteria, the Cytophaga–Flavobacteria–Bacteroidetes (CFB) group, and Alphaproteobacteria, with the aggregate-associated communities being the most distinct. At subsurface stations in the high-Arctic fjord Kongsfjorden, Jain et al. (2019) found that PA communities were enriched in carbohydrate-degrading Bacteroidetes (and Verrucomicrobia), consistent with the general association of Bacteroidetes with particles.

However, as shown in Table 2, our results were not consistent with results from those studies. For example, the relative abundance of Alphaproteobacteria was higher in 3 and 20 µm filters than in 0.2 µm. Also, Gammaproteobacteria was highly abundant in 0.2 µm filter (40.4 %). On the other hand, Bacteroidia showed higher abundance in 3 µm filter than other filters. Even if Alphaproteobacteria tend to be found in the FL fraction and Gammaproteobacteria and Bacteroidia in the PA fraction, the degree of FL–PA separation is context-dependent and strongly shaped by hydrography. Moreover, salinity gradients, suspended-particle load, and water residence time can outweigh a simple FL vs. PA dichotomy (LaMontagne & Holden, 2003; Selje & Simon, 2003; Zhang et al., 2007), and in some particle-rich estuaries the two fractions are almost indistinguishable due to rapid exchange between them (Hollibaugh et al., 2000). This probably applies to our study area. The Bering and Chukchi shelves are shallow, and the water column was weakly stratified in June 2017. Under these conditions, cells can move between the free-living and the particle-associated state. This kind of exchange would make the two fractions look more alike, not less, so the separation we found is best read as a minimum. It may also help explain why the environmental signal was weak here, with salinity the only significant variable (adjusted R² = 0.10). Therefore, in the Bering and Chukchi regions, which are undergoing rapid environmental change, the relative abundances of these bacterial classes may not always follow the typical pattern of dominant classes observed in FL and PA fractions.

### 4.2 Microbial Community by Depth

In temperate and other lower-latitude systems, depth is a consistent factor of structuring FL and PA bacterial communities, with community richness generally increasing with depth and a shift from surface oligotrophic FL groups to PA and deep water clades. In the northwestern Mediterranean, depth explained about 30 % of community variation (Crespo et al., 2013), and in the eastern Mediterranean FL diversity increased with depth as SAR406 and SAR202 became more abundant in deeper water (Roth Rosenberg et al., 2021). In the California Current, SAR11, Thaumarchaeota, and Rhizaria increased with depth, while surface phytoplankton-associated groups declined (Valencia et al., 2022), and in the Mauritania upwelling system, Bacteroidetes decreased and Deltaproteobacteria increased with depth, with FL–PA dissimilarity rising toward 200 m (Bachmann et al., 2018). In the open ocean, large sinking particles carried surface bacteria down into the deep sea, so that surface and deep communities resembled each other, whereas freely drifting (free-living) cells did not sink and their communities differed with depth (Mestre et al., 2018). On fast-sinking particles, the number of bacterial taxa decreased sharply with depth, until only a few fast-growing and nutrient-loving (copiotrophic) taxa remained (Baumas et al., 2021). Even in the Southern Ocean, depth was significant, but it was less important than water masses and size fractions (Milici et al., 2017).

In the Arctic Ocean, community composition also changes with depth, as in temperate regions, even though the shallow shelves cover only a small range of depths. In the Chukchi Borderland, depth was the dominant factor of community variation. Alphaproteobacteria, including SAR11, decreased sharply with depth while Deltaproteobacteria increased, and depth alone explained more than 80 % of their variance (Han et al., 2015). Community richness increased with depth, and assemblages separated between the surface/subsurface-chlorophyll-maximum layer and deeper waters in the central Arctic (Balmonte et al., 2018) and in Fram Strait, where PA communities became progressively enriched in Gammaproteobacteria, Bacteroidia, and Planctomycetes with depth (Fadeev et al., 2021). In the high Arctic fjord Kongsfjorden, particle-associated diversity was higher and community assembly became more strongly driven by stochastic processes at greater depth (Jain et al., 2021). Nutrient and particle characteristics also change markedly with depth in Arctic shelf and coastal waters (Kellogg & Deming, 2009, 2014).

In our study, ASV richness did not show a correlation with depth (Figure 9). Also, the correlation between ASVs in each size fraction (20, 3, and 0.2 µm) and depth (surface, middle, and bottom) was not statistically significant (Fisher’s exact test, p = 0.95). This is not consistent with the changes in community composition with depth, commonly reported in deeper Arctic basins and open-ocean settings, and most likely reflects the shallowness of the Bering and Chukchi shelves, with a well-mixed or only weakly stratified water column (Figure 14 in Shibue et al., 2026). Roth Rosenberg et al. (2021) reported that lifestyle such as FL vs. PA can shape bacterial communities more strongly than depth or seasonality, with PA communities responding dynamically to bloom-derived organic matter as active degraders. Given this, the strong vertical gradients in light, temperature, oxygen, and organic matter supply that drive depth-related community change in deep basins might be weaker in this region during June 2017. Although depth commonly influences the Arctic bacterial communities, in the Bering and Chukchi Seas, depth might become a less important factor shaping bacterial communities when the stratification is weakened due to a temperature anomaly.

### 4.3 Associations With Physicochemical Parameters

In temperate regions, salinity and particle concentration are the physicochemical variables most consistently linked to FL and PA bacterial community composition, together with temperature and phytoplankton-derived organic matter.

Along estuarine and coastal salinity gradients, salinity is the primary control. In the Baltic Sea, Gammapro-teobacteria increased and Actinobacteria and Betaproteobacteria decreased with salinity, while PA Bacteroidetes increased toward fresher water (Rieck et al., 2015). In the Weser estuary the proportion of PA bacteria tracked turbidity and suspended matter (r² up to 0.95), and Cytophaga–Flavobacteria decreased from the freshwater to the marine section (Selje & Simon, 2003).

Particulate loads and the quality of organic matter are also important for the PA fraction more generally. In Victoria Harbour, suspended solids best explained the differences between communities (Zhang et al., 2007). During a North Sea bloom, combined carbohydrates and POC shaped the PA communities, and Gammaproteobacteria were related to carbohydrates, POC, chlorophyll a, and temperature, while Bacteroidetes were related to chlorophyll a (Sperling et al., 2017).

Temperature often acts as a water mass proxy. In the Mauritania upwelling system, temperature was the best predictor of the FL community, whereas the PA community was explained by temperature together with ammonium (Bachmann et al., 2018). By contrast, the FL fraction tended to correlate with inorganic nutrients and phytoplankton pigments (Roth Rosenberg et al., 2021). Interestingly, some open ocean and deep sea studies did not link specific taxa to specific environmental factors. Rather than physicochemical variables, they described community differences in terms of depth, particle size, or how fast particles sink (Baumas et al., 2021; Mestre et al., 2018).

In the Arctic, the same variables matter, but freshwater input and water masses play a much bigger role. Salinity is the variable most closely linked to microbial communities. Also, salinity serves as an indicator of dissolved organic matter from these sources as it is freshened by terrestrial or glacial fresh water. Along the Mackenzie–Beaufort gradient, salinity shaped community composition and changed together with DOC, CDOM, and silicate, and using canonical correspondence analysis (CCA), open sea Alphaproteobacteria were separated from riverine Betaproteobacteria (Ortega-Retuerta et al., 2013). In the Chukchi Borderland, Alphaproteobacteria, including SAR11, correlated negatively with salinity, temperature, and density and positively with dissolved oxygen (Han et al., 2015), and temperature, salinity, and oxygen jointly explained surface-to-deep variation in the central Arctic (Balmonte et al., 2018).

River and glacial freshwater also play an important role, as the sources of nutrients and terrestrial organic matter to the Arctic Ocean. In the Laptev Sea, silicate and fresher water were associated with aggregate communities (Kellogg & Deming, 2009). In Svalbard fjords, DOC, nutrients, and *δ*^13^C followed meltwater and seasonal changes (Delpech et al., 2021; Jain & Krishnan, 2017). As in temperate waters, the FL fraction correlated with inorganic nutrients such as nitrite, phosphate, and silicate, whereas the PA fraction correlated with particle chemistry, including particulate carbohydrate, *δ*^13^C, C:N ratio, and dissolved oxygen (Jain et al., 2019). In the North Water and Beaufort Sea, phosphate and phytoplankton biomass (chlorophyll a and phaeopigment) correlated with all fractions (Kellogg & Deming, 2014). In the western Arctic, carbon sinking flux strongly correlated with chlorophyll a (r = 0.96) and negatively with the C:N ratio, and a diatom- and Gammaproteobacteria-rich group correlated with salinity (Qin et al., 2025). A time-series study in Fram Strait reported that bacterial groups such as Flavobacteriales correlated with sea-ice concentration and particle fluxes (Cardozo-Mino et al., 2023).

In this study, salinity was the only environmental variable with a significant marginal effect in the dbRDA (F = 3.15, p = 0.004, Figure 12). Though the overall model explained only a modest fraction of the variation (adj. R² = 0.10), it is consistent with previous studies (Han et al., 2015; Ortega-Retuerta et al., 2013) in which salinity is the dominant environmental factor of bacterial community composition. Jain et al. (2021) found that salinity explained approximately half of the variation in both FL and PA communities with a dbRDA in a high-Arctic fjord. Therefore, among the environmental variables, the significance of salinity in the dbRDA reinforces the interpretation that water mass structure, rather than depth alone, can be the primary factor distinguishing bacterial communities on the Bering and Chukchi shelf.

This idea helps us understand how each cluster was separated. For example, Clusters PA1 and PA3 showed substantially similar community compositions (lowest mean Bray–Curtis dissimilarity, 0.562) and had almost identical class-level compositions, both dominated by Bacteroidia (42.4 and 47.7 %, respectively) with comparable proportions (Bacteroidia to Gammaproteobacteria to Alphaproteobacteria was 5:3:2). The separation between Clusters PA1 and PA3 was driven by environmental factors rather than by broad differences in community composition. Cluster PA3 was found in deeper and more saline water than Cluster PA1 (median depth 25.9 vs 16.0 m; salinity 32.5 vs 31.9) and had higher NH_4_ (1.48 vs 0.31 µM) and PO_4_ (0.96 vs 0.73 µM), but lower NO_3_ (0.60 vs 1.30 µM). Therefore, Cluster PA3 occupied a deeper, more saline, ammonium-rich water mass, whereas Cluster PA1 corresponded to a shallower, fresher, nitrate-bearing but ammonium-poor water. Although Cluster PA3 occurred in deeper water, this separation does not necessarily mean that depth itself distinguished the communities in this region. Instead, water mass properties that correlate with depth, especially salinity and nutrient concentrations, can likely be the factors. Additionally, it is worth noting that NH_4_ is often interpreted as a remineralization product. However, since Cluster PA3 had many deep samples, high concentration of NH_4_ could also have been derived from benthic (sediment) sources. Moreover, its bloom index value was 0.46 (Figure 11), meaning that it could not be classified as a strong pre-bloom cluster and may possess both pre- and post-bloom characteristics. The two datasets differed in which environmental variables significantly structured the communities. While all five environmental variables were significant in the survey (five variables, p < 0.008), salinity was the only significant variable in the process dataset (p = 0.004). This contrast may partly reflect the influence of size fractionation, which likely accounts for a large share of the community’s variation in the process dataset and may have weakened the environmental effects that could be detected (adjusted R² *≈* 0.10). The process dataset also covered a smaller area with fewer samples than the survey dataset, so temperature, dissolved oxygen, and nutrients varied less, and they might correlate less clearly with the community composition.

## Data availability

Raw 16S rRNA gene amplicon reads from the ASGARD 2017 samples are available in the NCBI Sequence Read Archive under BioProject PRJNA421293 (SRA study SRP126234; https://www.ncbi.nlm.nih.gov/bioproject/PRJNA421293). All R scripts and outputs are publicly available at https://github.com/ReiSHIBUE/ASGARD2017_analysis.

## Author contributions

R.S. analyzed the data and wrote the manuscript. R.E.C. supervised the study.

## Acknowledgements

We thank Rachel M. Lekanoff, who participated in the ASGARD 2017 cruise, collected the DNA and environmental samples, processed them at the UAF Water and Environmental Research Center and the UAF DNA Core Lab, and provided the microbiological, hydrographic and environmental dataset.

## Funding

This work was supported by Genome Canada, the Natural Sciences and Engineering Research Council of Canada (NSERC) Discovery Grant Program (RGPIN-2022-05280 to R.E.C.), and the Canada Research Chairs Program (R.E.C.).

## Competing interests

The authors declare no competing interests.

## Footnotes

This manuscript is adapted from Chapter 3 of R. Shibue’s MSc thesis, *Marine Microbial Communities in a Rapidly Changing Arctic* (University of Manitoba, 2026).

## Notes

https://www.ncbi.nlm.nih.gov/bioproject/PRJNA421293

https://github.com/ReiSHIBUE/ASGARD2017_analysis

### Competing Interest Statement

The authors have declared no competing interest.

